# Changing the Fate of Dystrophin-Deficient Myoblasts via Hetero Ligand Nanoclusters on Biomaterial Surface: Effects of Integrin-Syndecan or Dystroglycan Crosstalk

**DOI:** 10.64898/2026.04.13.717576

**Authors:** Shirin Nour, Kristy Swiderski, Annabel Chee, Kate T. Murphy, Kevin I. Watt, Paul Gregorevic, Chayla L. Reevez, Amy Gelmi, Gordon S. Lynch, Andrea J. O’Connor, Greg Qiao, Daniel E. Heath

**Affiliations:** Department of Biomedical Engineering, Graeme Clark Institute, The University of Melbourne, VIC 3010, Australia; Polymer Science Group, Department of Chemical Engineering, The University of Melbourne, VIC 3010, Australia; Centre for Muscle Research, Department of Anatomy and Physiology, The University of Melbourne, VIC 3010, Australia; Novo Nordisk Foundation Centre for Stem Cell Medicine, Murdoch Children’s Research Institute, Royal Children’s Hospital, Parkville, VIC 3052, Australia; STEM College, School of Science, RMIT University, Australia

**Author notes:** Corresponding author: Associate Professor Daniel Heath.

**Keywords:** Biointerface, dystrophin-deficiency, mechanotransduction, syndecan, dystroglycan, tissue regeneration, neuromuscular co-culture

## Abstract

Engineering skeletal muscle tissue regeneration, particularly in dystrophin-deficient muscles is dependent on facilitating myogenesis and recovery of myotube structure and function, which can be challenging due to compromised cell-extracellular matrix (ECM) interactions. The current study explored the potential impact of enhancing dystrophin-associated protein complex and focal adhesion formation and the interaction with associated target receptors to improve cellular response in both normal and Duchenne muscular dystrophy (*Dmd*) mutant myoblasts. This was achieved by multivalent dual ligands functionalization of RAFT-synthesized copolymer with fibronectin- and laminin-derived adhesion peptides (RGD, AG73, and A2G80) and their clustering at the biointerface. Our findings demonstrated the synergistic effect of integrin-syndecan/dystroglycan engagement and their clustering on enhancing myoblast adhesion, proliferation, and differentiation, partially overcoming the deficits caused by loss of dystrophin. Furthermore, enhanced focal adhesion formation and elevated receptor localization, particularly dystroglycan, at the sarcolemma were associated with improved structural organization, mechanical stability, and neuromuscular connectivity of myotubes. These results suggest a novel insight into harnessing next-generation molecularly engineered biomaterials with robust interaction with cells’ mechanosensors for advancing skeletal muscle tissue engineering, offering potential applications in the regeneration of dystrophic muscle and the development of neuromuscular disease models for drug testing.

Graphical Abstract/ToC
Current work developed molecularly engineered biomaterial surfaces with nanoscale clustering of integrin-, syndecan-, and/or dystroglycan-binding peptides for skeletal muscle tissue regeneration. By controlling peptide distribution and type at the biointerface, cell adhesion, proliferation, and differentiation were modulated in dystrophin-deficient myoblasts. Accordingly, the results demonstrated significant improvement in myotube structural organization, mechanical stiffness, and their innervation in response to heteronanoclusters.

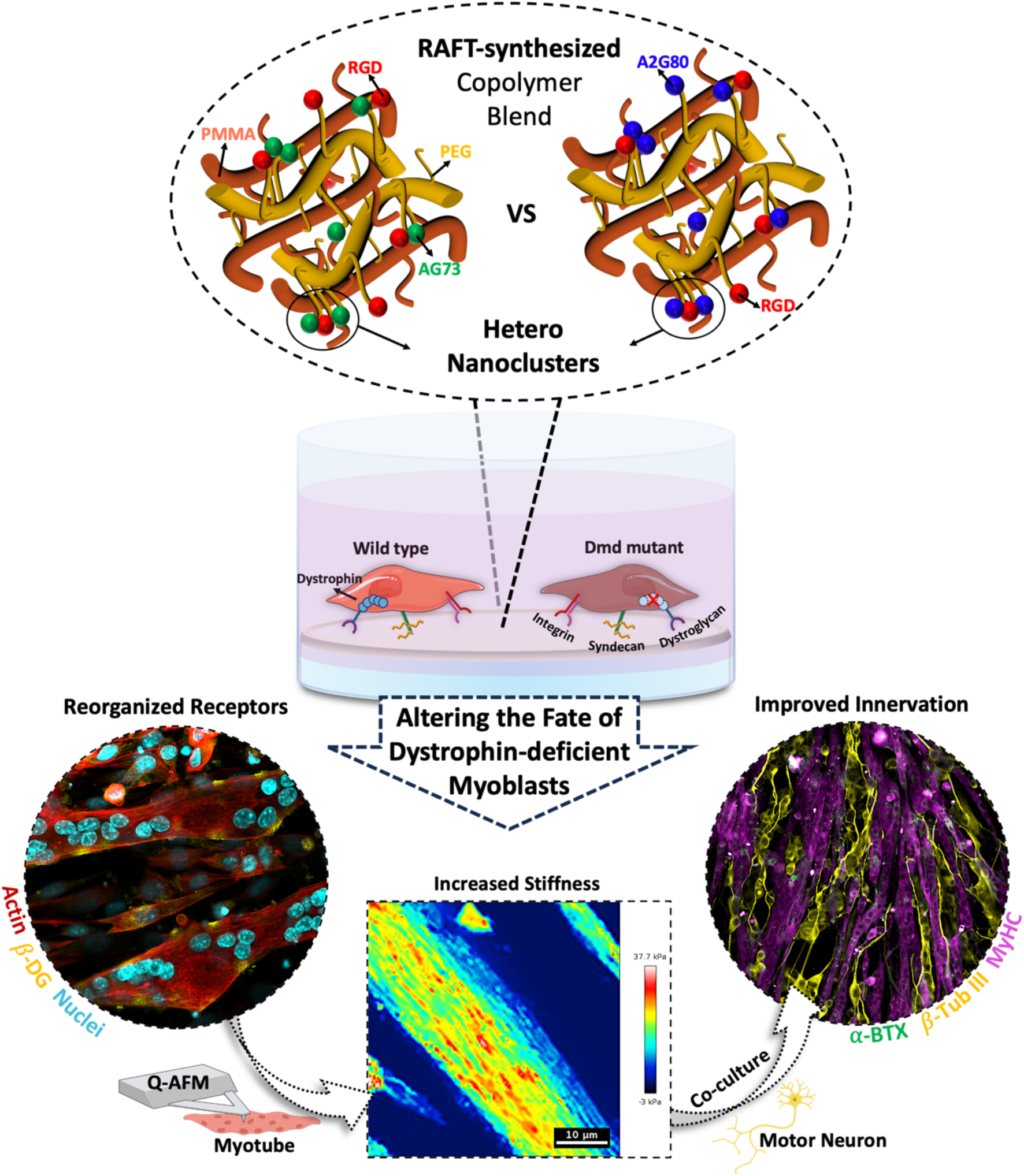

## 1. Introduction

Normal function and homeostasis of skeletal muscle tissue necessitate robust and dynamic communication between cells and their microenvironment through a myriad of transmembrane receptor binding events and intracellular protein complexations. In skeletal muscle cells, anchorage of actin cytoskeleton to the extracellular matrix (ECM) is not only accomplished by focal adhesion (FA)-mediated signal transduction but also by the parallel assembly of integral and peripheral membrane proteins known as the dystrophin-associated protein complex (DAPC).

Integrin receptors (e.g., α_5_β_1_ and α_3_β_1_) are the primary cell-ECM adhesion receptors binding specific polypeptide motifs or ligands (including RGD tripeptide from fibronectin). While integrin-ligand interactions are essential, recent studies revealed the crucial role of co-receptors such as syndecan-4 (a ubiquitous member of heparan sulphate proteoglycans), in mechanotransduction and maturation of FA. In addition to interacting with laminin and growth factors within the ECM for cell adhesion, the cytoplasmic domain of syndecan-4 plays as key role in the regulation of proliferation and migration of myoblasts via PKCα activation and RhoA signaling ^[1–3]^.

On the other hand, dystrophin acts as a mechanotransducer and mechanosensor through interaction with dystroglycan as one of the key glycoproteins in the DAPC, which also has crosstalk with the integrin complex ^[4,5]^. Dystroglycan is comprised of two structurally connected subunits, including the extracellular α domain, as a laminin-binding adhesion molecule, and a transmembrane β domain, which binds the α domain and dystrophin to confer mechanical stability to myotubes during the continuous cycles of contraction and relaxation ^[6,7]^.

Dystroglycan also contributes to the stabilisation and clustering of acetylcholine receptors ^[8,9]^. Moreover, there is evidence that syndecans are involved in signalling pathways related to astrocyte polarity, axonal guidance, neurite outgrowth, and regulation of synapse development. These are suggestive of the crucial role of syndecan and dystroglycan in post-synaptic organization during neuromuscular junction (NMJ) formation ^[8,10–12]^.

Dystrophin deficiency, such as that occurring in muscular dystrophies and neuromuscular degenerative diseases, leads to DAPC destabilization, mislocalization/degradation of β-dystroglycan in sarcolemma, impairing the structural integrity of muscle and negatively affects FA and homeostatic cytoskeletal tension. Despite the determinant role of DAPC and FA in maintaining normal skeletal muscle cell behavior and their functional overlap, the mutual interaction/crosstalk between involved receptors and their impact on muscle tissue regeneration, particularly in dystrophic conditions, remains unexplored ^[5]^.

One of the promising approaches in controlling cellular behavior and fate is via the development of advanced biomaterials mimicking the native microenvironmental cues at the biointerface to modulate adhesion receptor binding ^[13]^. We have recently shown that effective engagement of integrin using nanoscale clustering of integrin binding peptides at the biomaterial surface can improve myoblast adhesion, growth, and differentiation controlled by density and presentation of the adhesion ligands ^[14–17]^. We and others also demonstrated the positive impact of the biomaterials systems with syndecan-binding peptides on cell adhesion, motility, shape organization, and self-renewal ^[1,15,18]^.

Building on our previous works of peptide-functionalized RAFT synthesized polymer ^[14]^, we developed the heterogenous form of multivalent peptide nanoclusters by combining integrin-binding ligand (RGD) with syndecan-4- or dystroglycan-binding peptides (AG73 and A2G80, respectively). Then, we investigated the behavior of C2C12 myoblasts in normal (wild type) and dystrophin-mutant conditions on the fabricated substrates and their ability to form NMJ. We hypothesized that supporting clustering of integrin-syndecan or integrin-dystroglycan within the cell membrane would contribute to reinforcing FA and stabilizing DAPC components, including β-dystroglycan, to improve cytoskeletal organization and subsequent cell function even in dystrophin-deficient cells.

The results suggested that our bioengineered materials further enhanced cell spreading, focal adhesion formation, proliferation, and differentiation compared to control and homocluster surfaces. Interestingly, analysis of the myotube’s structural organization and stiffness revealed significant improvements in the mechanical stability of dystrophic myotubes along with β-Dystroglycan clustering within the membrane which has not been reported before. Furthermore, compared to RGD nanoclusters, the heterocluster of peptides increased neuromuscular interactions examined with acetylcholine receptor (AChR) clustering and neural branching even in dystrophin-deficient group. Our results, highlight the role of modulation of FA and DAPC formation and crosstalk in changing the fate of dystrophic skeletal muscle cells. This study has offered a promising biomaterial engineering approach for possible fundamental studies on muscle regeneration and drug screening applications.

## 2. Results and discussion

### 2.1. Fabricating cell culture surfaces with nanoclusters of integrin-, syndecan- and dystroglycan-binding peptides

To prepare the peptide nanoclustered surfaces, two amphiphilic random comb polymers were synthesized via reversible addition-fragmentation chain-transfer (RAFT) controlled radical polymerization based on a previously developed protocol ^[14,15]^. Specifically, a non-functionalized copolymer of methyl methacrylate and poly(ethylene glycol) methacrylate (hereafter referred to as the MP polymer) and terpolymer of methyl methacrylate, poly(ethylene glycol) methacrylate, and hydroxyl terminated-poly(ethylene glycol) methacrylate (hereafter referred to as the MPP polymer) were synthesized with molecular weights between 120-170 kDa and low PDI (1.2-1.3). The hydroxyl-terminated pendant groups of the MPP polymer were functionalized with norbornene groups via an esterification reaction. Thiol-ene click chemistry was then employed to attach cysteine-terminated peptides to the polymer chains (**Figure 1A**). The quantity of peptide in each functionalized polymer was assessed via trace elemental analysis.

**Figure 1.**
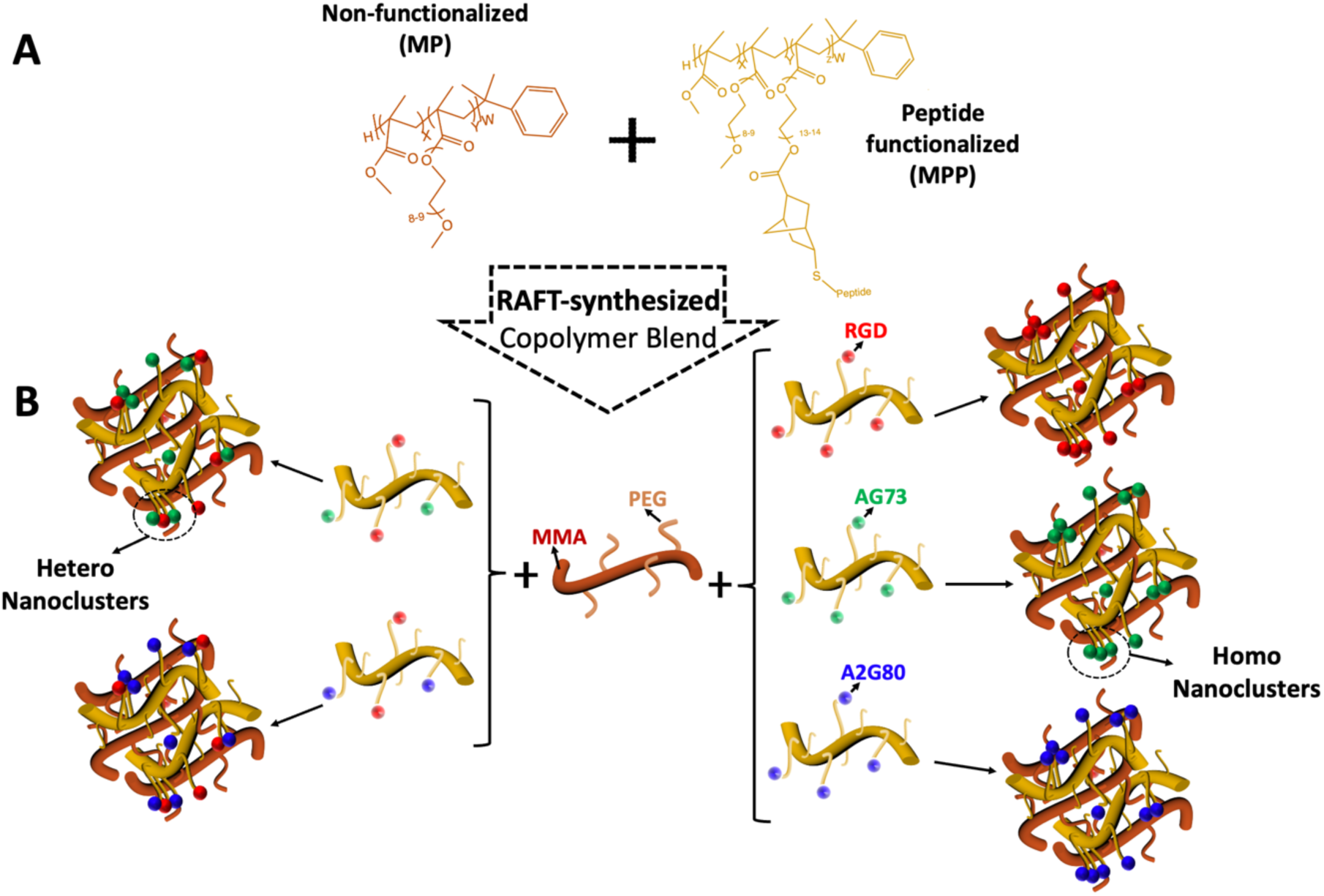
A) Chemical structure of the RAFT synthesized polymers including non-functionalized (MP) and peptide-functionalized (MPP) polymers. B) Schematic illustration of peptide nanocluster surfaces formed by blending of MP copolymer (orange) and MPP terpolymer (yellow) functionalized with RGD (red), AG73 (green), A2G80 (blue) to form homocluster surfaces or a combination of RGD with AG73/A2G80 to form heteroclusters.

The percentage of PEG-pendant groups in the polymer was held at approximately 15 mol% because earlier studies by us and others have shown that these comb polymers can be fabricated into water-stable cell culture surfaces with excellent resistance to protein fouling and a thermodynamic driving force (autophobicity) that enriches peptides at the interface ^[14,15,19,20]^. Full details on the polymer synthesis and characterization by ^1^H NMR and GPC can be found in the Supplementary Information (**Figure S1-3 and Table S1-3**).

Surfaces containing nanoscale clusters of integrin-, syndecan-, and dystroglycan-binding ligands were created by blending specific ratios of MP and MPP polymers and casting them info films. This blending technique allows us to control the global ligand density (μg peptide/mg polymer) by controlling the ratio of functionalized to non-functionalized polymer chains and the local ligand density by controlling the number of ligands per functionalized chain. ^[14,15,20]^. This technique allows us to independently control the average ligand densities while molecularly patterning the ligands into a nanoscale cluster.

Previously, we have demonstrated that cell culture surfaces with a local ligand density of 4 peptides per MPP polymer chain and a global ligand density of 7 ug of RGD peptide/mg polymer was optimal for the skeletal muscle development, resulting on more myotubes that were also more mature ^[14]^. However, syndecan-4 and dystroglycan receptors are also key in directing skeletal muscle development and phenotype. Therefore, we developed cell culture surfaces that also displayed peptides that bind these other receptor types, as detailed in **Table 1** (peptide synthesis and purification details are provided in the **Supplementary Information, Figure S1**). The RGD peptide was selected as the most frequently used integrin-binding ligand due to its strong affinity for focal adhesion receptors α_3_ and α_5_β_1_. To target syndecan-4 and dystroglycan, AG73 and A2G80 were selected, respectively, based on previous studies ^[15,21–23]^. Additionally, as negative control surfaces, non-functionalized polymer and polymer functionalized with the non-adhesive RGE peptide and scrambled versions of the syndecan-and dystroglycan-binding peptides were used.

**Table 1.**
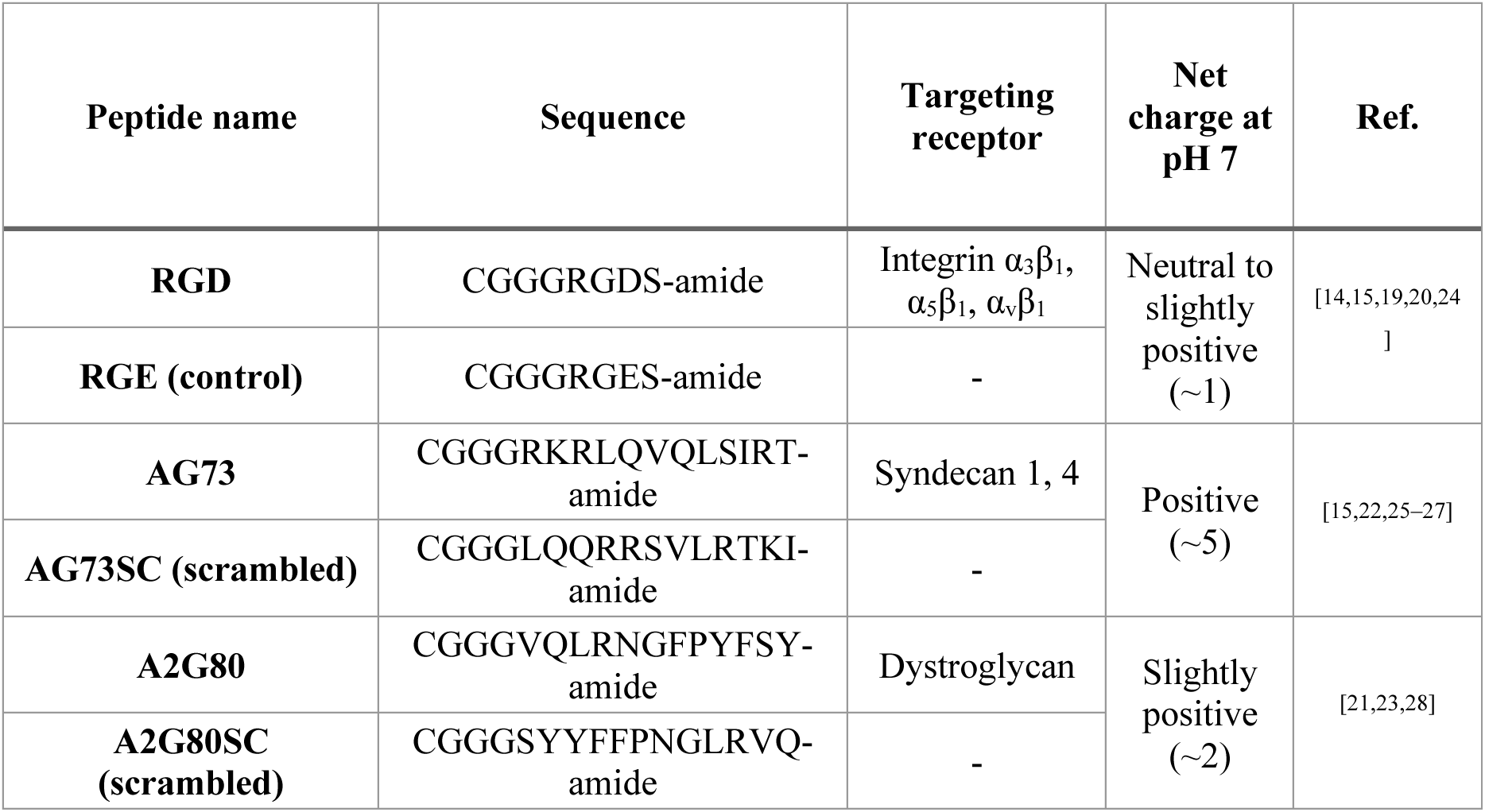
Cell adhesive peptides used in the fabrication of cell culture surface, their amino acid sequence, known receptor binding behaviour, and net charge at pH 7.

The polymer blending technique also enables control over the mixing of ligands in each nanocluster, as shown in **Figure 1B**. Specifically, if MPP polymer chains are functionalized with only one ligand type, films with “homoclusters” of ligands are produced. In contrast, if MPP polymer chains are functionalized with multiple ligand types, “heteroclusters” of ligands are produced. In this work, cell culture surfaces with homoclusters of the RGD-, AG73-, and A2G80-peptides and heteroclusters of RGD and AG73 or RGD and A2G80 were produced (**Figure 1B**) and blending with a non-functionalized polymer. In the case of heterocluster surfaces, different mol ratios of RGD:AG73/A2G80 were used (70:30, 50:50, 30:70). Additionally, a randomly functionalized RGD surface (rRGD) (without blending) was included as a control. The various surfaces used in this work are compiled in **Table 2**.

**Table 2.**
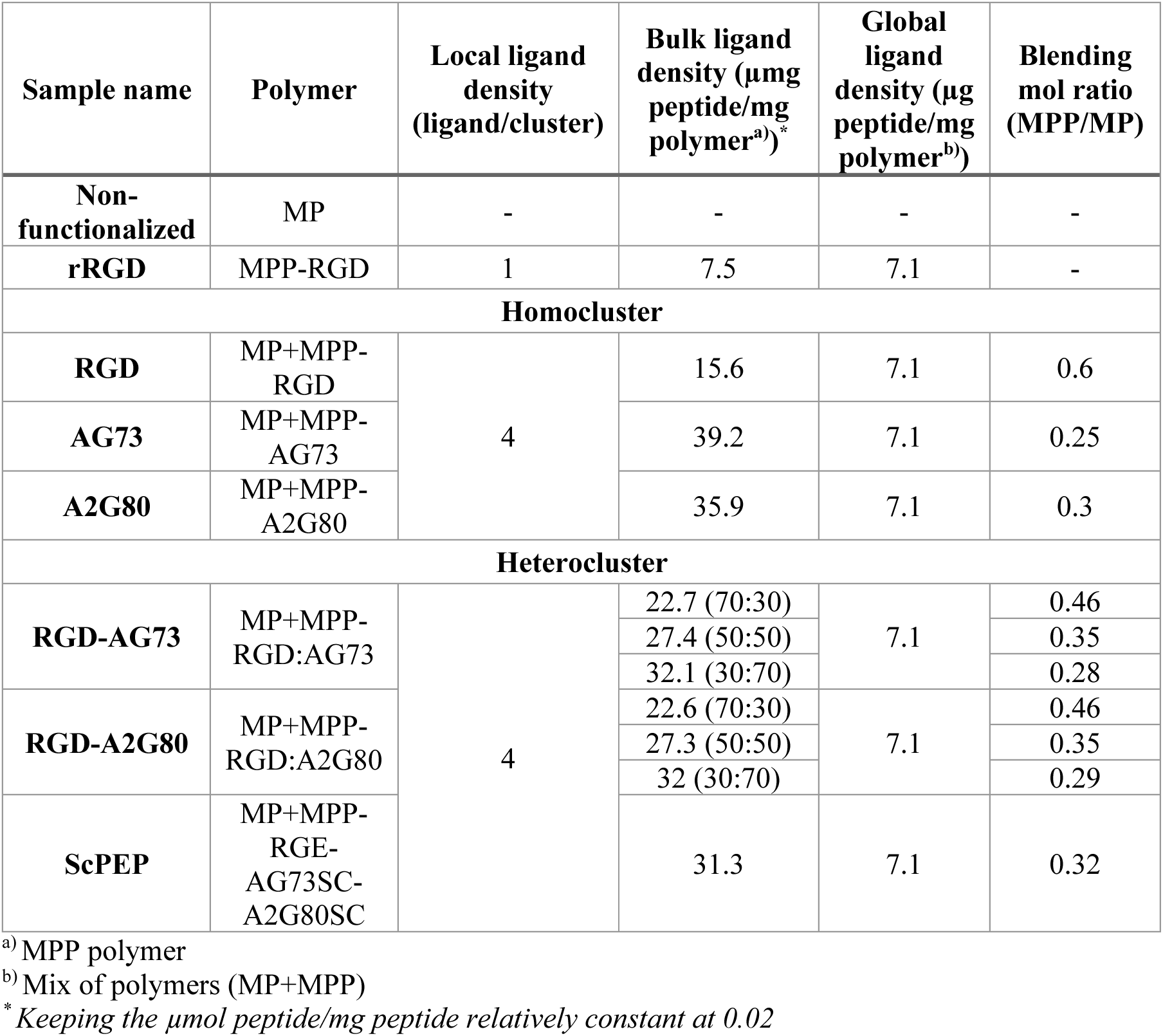
Polymer surfaces used in this study.

### 2.2. Physicochemical characterization of peptide conjugated polymer surfaces: formation of nanoclusters of ligands with surface potential depending on ligand type

We further characterized our developed polymer surfaces using Raman and EDX spectroscopies and KPAFM technique to analyze the presence and distribution of peptides on the surface depending on the ligand type (**Figure 2**). Accordingly, as shown in **Figure 2A**, the Raman spectra in all groups displayed characteristic peaks of the polymer, including C-C bending and C-O stretching (815, 985 cm^−1^), CH_2_/CH_3_ bending (1445 cm^−1^), C=O stretching (1726 cm^−1^), and C-H and O-H stretching (2800-3100 cm^−1^). Noticeably, upon peptide conjugation, the emergence of a new peak attributed to the stretching of C-N (amide II) (1590 cm^−1^) is evident in peptide-functionalized polymers (a-c) compared to the non-functionalized polymer (d), confirming the presence of peptides.

**Figure 2.**
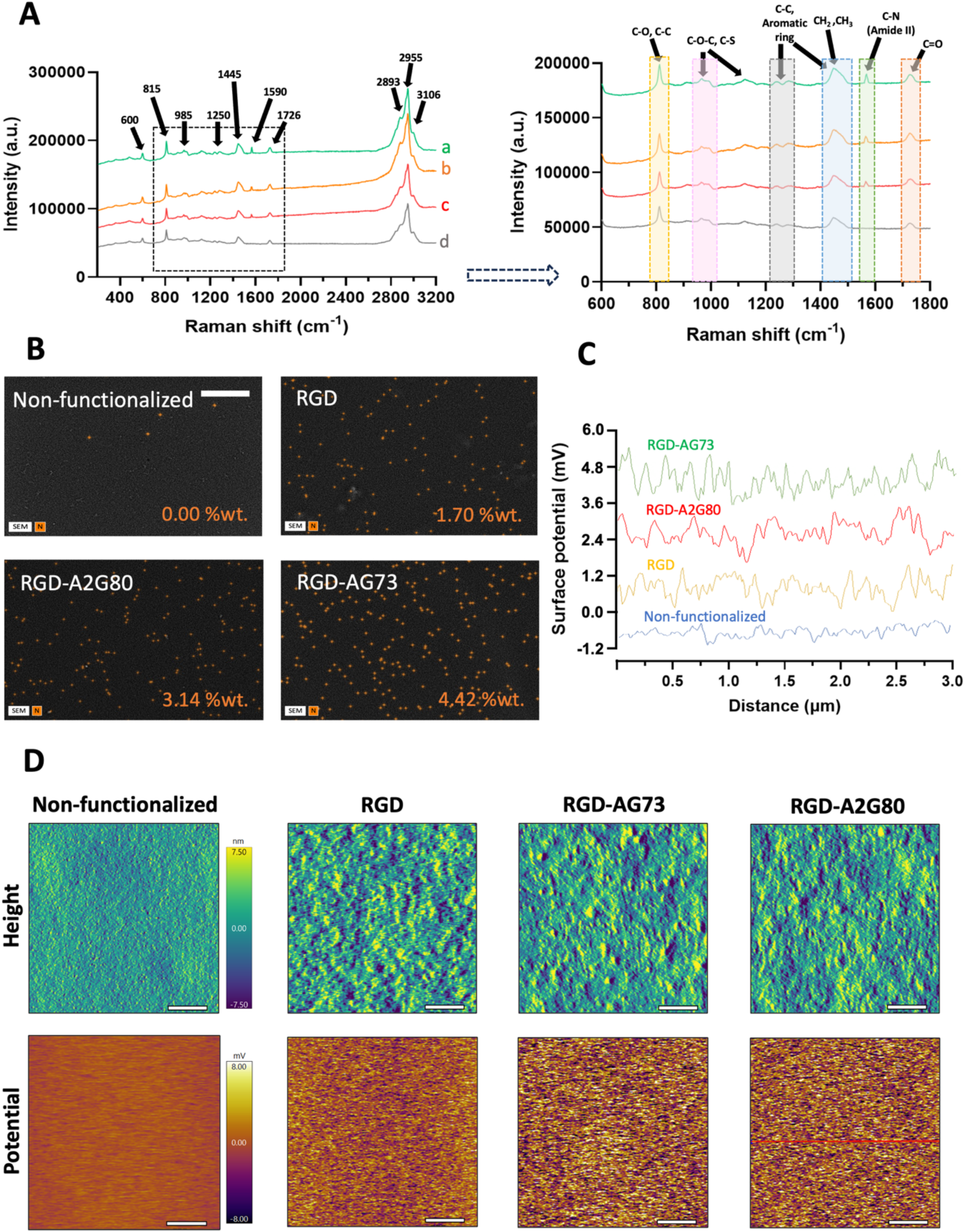
Evaluation of surface composition and peptide distribution. A) Raman spectroscopy supported the presence and covalent attachment of peptides to the functionalized polymer. B) EDX confirmed the presence of nitrogen related to the amine groups of the peptides. C, D) AFM analysis and images of the samples in KPFM mode which shows the surface topography (height map, in the top row of panel D) and charge distribution (potential map, bottom row panel D) across the polymer surfaces after equilibrium swelling in water. The calculated average surface potential illustrated with graph (C), which represents the distribution of surface charge across the middle of the sample (indicated with the red line in panel D image). Scale bars are 100 µm and 200 nm in panels B and D, respectively.

On the other hand, there was a noticeable increase in the peak intensity at 1250, 1445, 1590, and 1726 cm^−1^ in response to addition of peptides particularly, RGD-AG73 (b) and RGD-A2G80 (c) due to the longer peptide chain and more amine groups as well as presence of amide III and aromatic ring attributed to the proline, tyrosine and phenylalanine residues in A2G80. Moreover, the spectra of the peptide-functionalized polymers showed a slight shift and increase in peak intensity around 1100-1132 cm^−1,^ which can also represent the C-S vibration related to the peptides and changes in the glycol unit after conjugation. These results aligned with the NMR data in **Figure S3**, which confirmed successful conjugation of peptides to the polymer and their availability at the interface.

In addition, EDX analysis of the samples was utilized to assess the amount of atomic nitrogen and its distribution at the surface related to the amine groups found in the peptides. According to **Figure 2B**, as expected, no nitrogen was detected on the non-functionalized polymer in contrast to peptide-functionalized polymers, which confirmed the presence of nitrogen distributed across the interface. Despite constant local and global ligand density, moving from RGD to RGD-AG73 and RGD-A2G80, we observed slight increase in %wt. of nitrogen (approximately 1.8 and 2.6 times, respectively), which may be attributed to the difference in the nitrogen contents of AG73 (∼29) and A2G80 (∼22) peptides compared to RGD (∼12) (**Table 1** and **Figure S2**). Although the amount of peptide in each polymer sample and per cluster was the same, there were approximately 2-fold nitrogen atoms in AG73/A2G80, and considering the 50:50 ratio of functionalization with RGD and A2G80/AG73, the increased nitrogen %wt. in EDX results can be explained.

AFM in Kelvin Probe Force Microscopy (KPFM) mode was next performed on the samples to investigate the surface potential distribution related to the peptide functionalization of the polymer and their clustering. **Figure 2D** illustrates both the topographical image (top row) and the corresponding surface potential map (bottom row) of the scan area.

The non-functionalized polymer surfaces exhibited a relatively smooth surface with zero to slightly negative surface potential (**Figure 2C, D**). In contrast, after functionalization with peptides and blending, a change in surface topography and formation of nanoscale bumps were observed. We have previously shown that due to the quasi-2D molecular organization and autophobicity effect, at the interface of our polymer blend system, while the hydrophobic polymer backbone stays in random coil form facing the substrate, the hydrophilic peptide-functionalized PEG chains extend into the aqueous biological environment ^[29,30]^. This can lead to the observed surface feature in topography/height images.

On the other hand, potential map and quantitative analysis (**Figure 2C, D**) indicated a noticeable increase in surface potential, particularly in RGD-AG73, which may be attributed to the presence of positively charged amino acids such as lysine and arginine in the peptide sequence. The observed shifts in surface potential confirm successful peptide immobilization and their uniform distribution. The results demonstrate the sensitivity of KPFM in detecting subtle biofunctionalization on polymer surfaces. Understanding surface charge is crucial for the design and optimization of biointerfaces affecting cell adhesion.

### 2.3. Improved cell adhesion and proliferation: synergistic effect of syndecan-/dystroglycan-binding peptide with integrin binding peptide

To assess the impact of our biofunctionalization strategy and optimise the ratio between the ligands within heterogenous clusters, we first examined the cell morphology, focal adhesion formation, and density of normal C2C12 myoblasts. As seen in **Figures S4 and S5**, cells attachment was relatively low on both the non-functionalized polymer (MP) and the scrambled peptide functionalized sample (ScPEP) as controlled surfaces, as indicated by the low number and total area of focal adhesions (**Figure S4A-C**), which confirms the low fouling nature of the polymer and the selectivity of the peptides. In addition, it was apparent that ligand clustering significantly improved cell spreading and adhesion, with more than 2-fold increase in number and area of focal adhesions when cultured on cluster surfaces of RGD, RGD-A2G80, and RGD-AG73 with a 50:50 blending ratio compared to rRGD (**Figure S4A-D**). These results are in line with our previous observations on the effect of ligand clustering and density on myoblasts and endothelial cells’ adhesion and growth ^[14,15]^.

However, while myoblasts were able to elongate on the homogenous clusters of AG73 and A2G80 alone, as demonstrated by the vinculin (green) fluorescent signal, there was a noticeable decrease in the number and area of the focal adhesions relative to the RGD clusters (**Figure S4A-C**). This was expected due to the major role of integrin binding in the development of the focal adhesion complex. However, compared to control and the homogeneous RGD cluster, combining RGD with AG73 (RGD-AG73) or A2G80 (RGD-A2G80) promoted cell spreading and vinculin localization at adhesion sites, particularly at a ratio of 50:50 in RGD-AG73 (∼1.2-fold increase in focal adhesion area, **Figure S4A-D**). Moreover, after 3 days of culture, heterogeneous cluster surfaces comprised of RGD-AG73 and RGD-A2G80 with a 50:50 ratio improved cell proliferation compared to homogenous clusters and control, although this was more pronounced with RGD-AG73 (**Figure S4E**).

Based on this characterization, the 50:50 ratio was determined to be the optimized formulation of peptide heteroclusters, which was subsequently used to evaluate the impact of peptide clustering on the adhesion and morphology of dystrophin-deficient (*Dmd* mutant) myoblasts (refer to **Figure S6** for western blot verification of control and dystrophin-deficient myoblasts). Similar to previous observations, a significant increase in the number of focal adhesion and cell area was observed when cultured on heterogenous cluster surfaces compared to homogenous and control samples in both control and dystrophin deficient myoblasts (**Figure 3A-D**).

**Figure 3.**
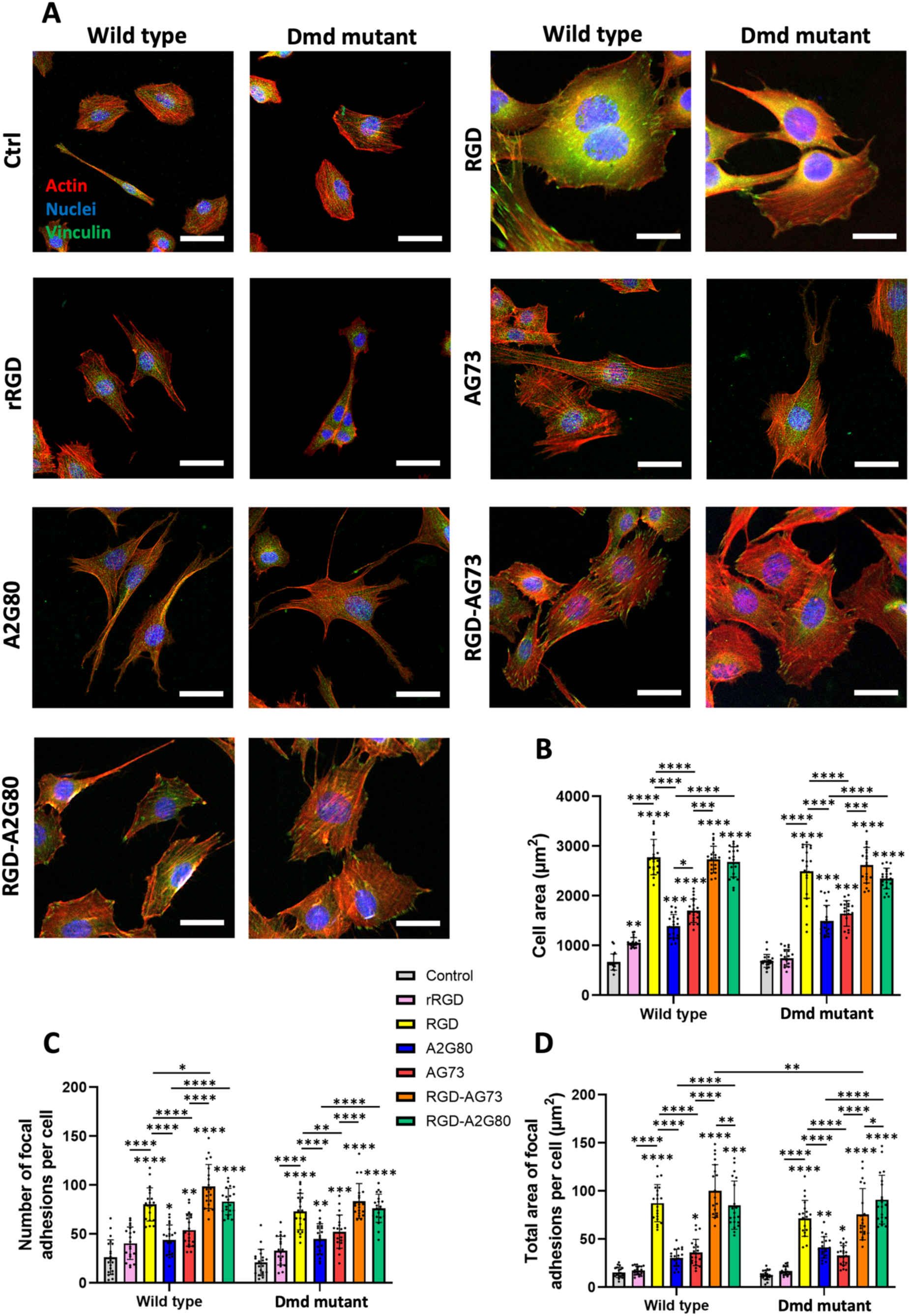
Representative confocal microscopy images of control (wild type) and dystrophin-deficient (*Dmd* mutant) C2C12 myoblasts after 1 day of culture on the polymer surfaces. A) Immunostaining against vinculin (green), F-actin (red), and nuclei (blue), Scale bar: 30 µm. B) Average cell areas as a function, C) Number of focal adhesions per cell, and D) Total area of focal adhesions per cell (n=20). Asterisks represent the statistical significance between the treatment groups, with ✻ directly above data points indicating statistical differences with the control, unless above the bars which indicate statistical differences between the treatment groups.

Additionally, considering the important role of neuromuscular connections on the function of engineered skeletal muscle tissue, we assessed the morphology, adhesion, and growth of NSC-34 motor neurons in response to the surfaces (**Figure S7**). Similarly, the positive synergistic effect of integrin binding peptides and syndecan/dystroglycan binding peptides on the degree of cell spreading, along with number and area of focal adhesion, was observed (**Figure S7A-D**). Moreover, compared to RGD clusters, heterogeneous peptides significantly altered the actin organization and enhanced the vinculin fluorescent signal (**Figure S7A, C**). In case of cell proliferation after 3 days, comparing the homogenous and heterogenous clusters, no obvious trend was detected, although A2G80 and AG73, performed slightly better than their heterogenous clusters version (**Figure S7E**).

These findings suggest that co-presentation of RGD with syndecan-binding peptide (AG73) or dystroglycan-binding peptide (A2G80) promotes synergistic crosstalk and related signalling pathways, facilitating stronger adhesion and accelerated cell proliferation, which may be useful for future biomaterials design for improved cell-material interactions. While it has been previously shown by both our group and others that integrin binding is crucial for cell adhesion, mature focal adhesion, and successful mechanotransduction-mediated signalling necessitates the engagement of co-receptors such as syndecan-4 ^[15,18,27,31]^. Syndecan-4 plays an important role in the recruitment of protein kinase Cα (PKCα) into the focal adhesion sites, initiating downstream signalling ^[32,33]^. Moreover, PKCα activation leads to the increased expression of RhoA signalling protein, which is crucial for the development of focal adhesions and actin stress fiber formation and requires cross-signalling between syndecan-4 and α_5_β_1_ integrin. In our designed biomaterial, while RGD served as an integrin binding ligand (interacting with α_5_β_1_ and α_3_β_1_), AG73 and A2G80 peptides targeted syndecan-4 and dystroglycan, respectively ^[21,23,25]^. Interestingly, although there is evidence of crosstalk between integrin and dystroglycan ^[34–37]^, this is one of the first studies to report a role for integrin-dystroglycan or syndecan in cytoskeletal organization and focal adhesion formation in dystrophin-deficient myoblasts.

### 2.4. Myogenesis and myotube structure is regulated by heterogeneity of ligand nanoclusters: enhanced differentiation, sarcomere organization, and myotube mechanical integrity

To assess the impact of ligand heterogeneity on the success of myogenesis, control and dystrophin-deficient C2C12 myoblasts were grown to confluence and differentiated into myotubes for 7 days. Myotube formation was evaluated via immunostaining for myosin heavy chain (MyHC) and sarcomeric ⍺-actinin as hallmarks of myotube formation ^[38]^.

Confocal microscopy images of myotube formation on polymer films under the impact of ligand type and heterogeneity, revealing significant differences in myotube morphology and density in both wild type and *Dmd* mutant cells (**Figure 4A**). Accordingly, while myotube formation was observed on all surfaces, the heterogeneous cluster surfaces generated longer and thicker myotubes compared to both the tissue culture plastic control (Ctrl) and homogeneous cluster surfaces, which showed sparse and thin myotubes (**Figure 4A**). This was reflected by an approximately 33% increase in fusion index (**Figure 4B**) and myotube diameter (**Figure 4C**) for control and dystrophin-deficient myotubes cultured on RGD-AG73 and RGD-A2G80 compared to control. In addition, there was a significant increase (approximately 1.8-fold increase) in the myotubes’ diameter on RGD-A2G80 compared to RGD alone.

**Figure 4.**
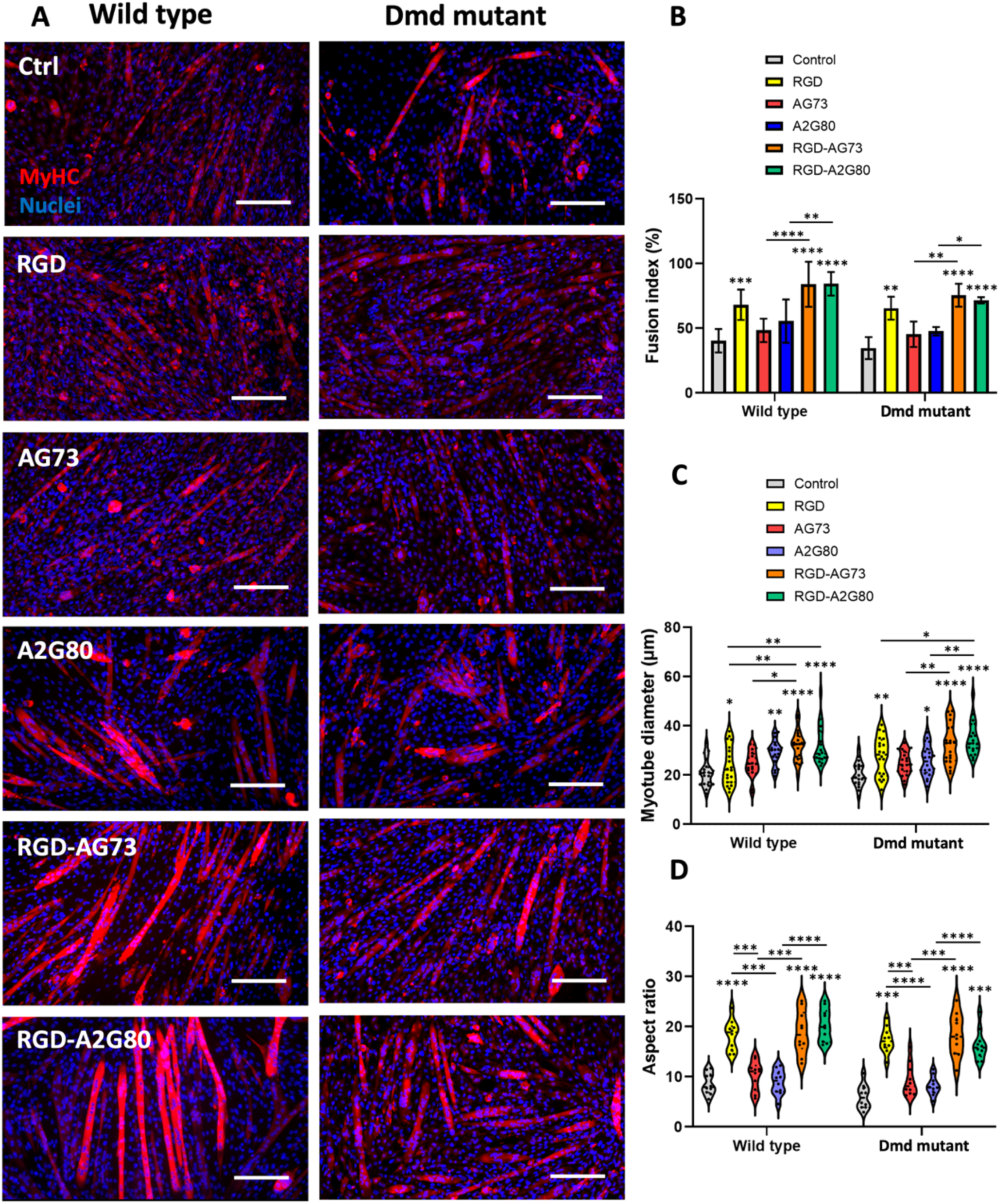
Evaluation of myotube formation on biomaterial surfaces. A) Confocal microscopy images of myotubes after 7 days of culture. Immunostaining against MyHC (red) and nuclei (blue), scale bar: 200 µm. Image analysis with ImageJ was performed to measure B) fusion index, C) myotube diameter, and D) aspect ratio (n=20). Asterisks represent the statistical significance between the treatment groups, with ✻ directly above data points indicating statistical differences with the control, and above the bars indicating statistical differences between the treatment groups.

On the other hand, based on **Figure 4D**, analysis of the myotubes’ aspect ratio (ratio of length to diameter) showed enhanced elongation of myotubes under the effect of peptide clustering in which while control and homogenous clusters (except RGD) exhibited the lowest aspect ratio, heteroclusters of RGD-AG73 and RGD-A2G80 conserved significantly longer tubes (higher aspect ratios) with no statistical difference between the wild type and *Dmd* mutant cells.

Further investigation of sarcomeric *α*-actinin staining of the myotubes (**Figure 5A, B**) revealed the compromised sarcomere formation in *Dmd* mutant cells compared to wild type, which was expected. On the other hand, provide peptide clustering significantly improved the sarcomere organization, particularly peptides heteroclusters, leading to highly striated myotubes with well-defined z-bands, interestingly in both *Dmd* mutant and wild type groups.

**Figure 5.**
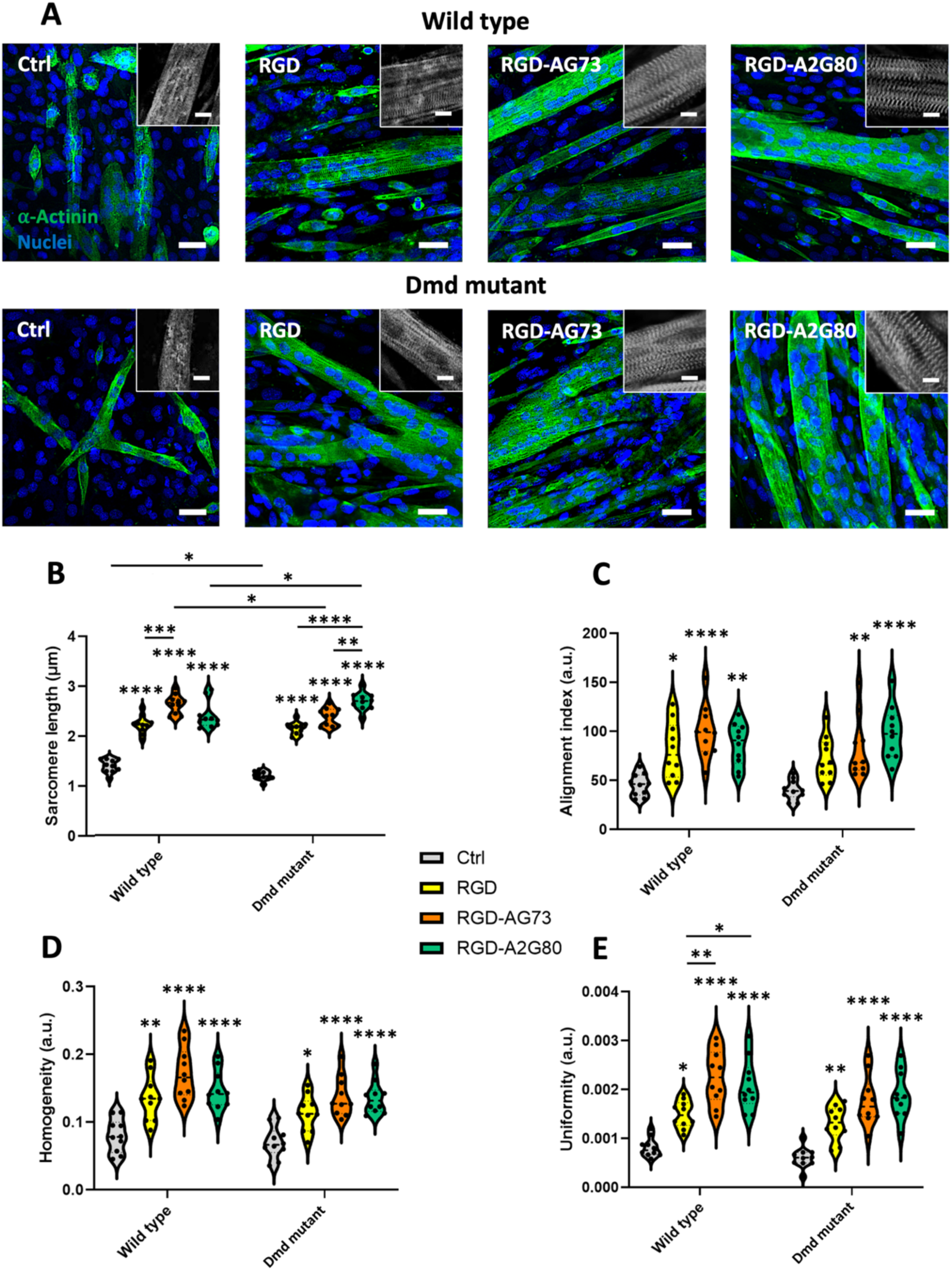
Analysis of sarcomere formation in response to peptide heteroclusters after 7 days. A) Confocal images of stained myotubes against sarcomeric ⍺-actinin (green), and nuclei (blue). Image analysis results using the Sota tool plugin of ImageJ software indicated an improvement in B) sarcomere length, C) alignment, D) homogeneity, and E) uniformity. Scale bars are 30 µm and 15 µm for zoom-in images (n=20). Asterisks represent the statistical significance between the treatment groups, with ✻ directly above data points indicating statistical differences with the control, unless above the bars which indicate statistical differences between the treatment groups.

Quantitative image analysis also confirmed these observations. Accordingly, as shown in **Figure 5B-E**, RGD-AG73 and RGD-A2G80 surfaces resulted in the highest sarcomere length and significantly higher alignment relative to the control. Notably, *Dmd* mutant cells treated particularly with RGD-A2G80 reached sarcomere lengths higher than wild type group with comparable structural homogeneity, alignment and uniformity (also observed in RGD-AG73), indicating a partial rescue of the defective phenotype. Similar improvement was also observed in case of sarcomere homogeneity and uniformity in response to heterocluster compared to control and RGD cluster. This suggests myotubes’ structural maturation and likely improved cytoskeletal tension and directional organization induced by effective engagement and possible clustering of integrin and its co-receptors, including syndecan and dystroglycan.

Furthermore, the ability of peptide’s heteroclusters, particularly RGD-AG73, to partially restore sarcomeric organization and uniformity in *Dmd* mutant cells compared to the control surface and wild type suggests that employing ECM-derived ligands can be effective to compensate for the cytoskeletal regulation caused by dystrophin deficiency. It has been shown that syndecan-4 is usually upregulated in dystrophic muscle cells ^[39]^. It is speculated that due to decreased dystrophin expression (**Figure S6**) and the possible disruption of mechanotransduction and mechanical instability, compensatory increases in syndecan-4 as the key focal adhesion co-receptor can occur ^[3,40]^. Therefore, the improved response we observed by culturing *Dmd* mutant cells on our heterocluster samples, particularly, RGD-AG73 functionalized biomaterial, reflects the robust engagement cell receptors and their targeting ligands, possibly activating FAK signalling and downstream pathways like ERK/MAPK.

On the other hand, the improved myogenesis and sarcomere formation under the effect of RGD-A2G80 in both wild type and *Dmd* mutant cells was unexpected, considering the decreased expression of dystrophin and dystroglycan in *Dmd* mutant cells (**Figure S5 and 6**). It has been shown previously that dystroglycan confers stability to myofibers during the continuous cycles of contraction and relaxation ^[7]^ by anchoring it to the ECM via DAPC formation. However, lack of dystrophin and DAPC destabilization may result in compromised dystroglycan (particularly the β subunit) localization in the sarcolemma and increase its degradation ^[41,42]^. This can disrupt the interaction between β-dystroglycan and the actin cytoskeleton, impairing the myotubes’ structural integrity and signalling functions of the DAPC and focal adhesion, which can further affect myotube stiffness and degradation over time ^[4,7,43]^.

We hypothesized that the observed positive effect on myogenesis and striation can not necessarily be related to the number of engaged dystroglycan-integrin receptors, but also a result of improved localization and clustering of the receptors within the myotube’s sarcolemma in response to our designed biomaterials. Therefore, to test our hypothesis, we evaluated the myotube’s mechanical stiffness using AFM nanoindentation and additional immunostaining against F-actin and β-dystroglycan.

Using live cell quantitative AFM (Q-AFM), detailed and real-time information about the structure and mechanical properties of myotubes, including stiffness through analysis of force-indentation distance curves, was obtained. As seen in **Figure 6**, culturing control myotubes on our peptide cluster surfaces, particularly heterogeneous clusters, significantly increased the measured Young’s modulus, with the highest stiffness achieved on RGD-A2G80 (∼32.4 kPa; **Figure 6A, B**). Notably, the Young’s modulus in dystrophin-deficient myotubes was significantly lower (∼43%) than that of control myotubes, which is likely explained by the lack of Dp427 dystrophin protein expression ^[35,44]^. Interestingly, stiffness of the dystrophin-deficient myotubes was similarly increased when cultured on the peptide clusters, increasing to approximately 21.4 kPa on the RGD-A2G80 surface; however, this remained lower than that of control myotubes on the same surface (**Figure 6B**).

**Figure 6.**
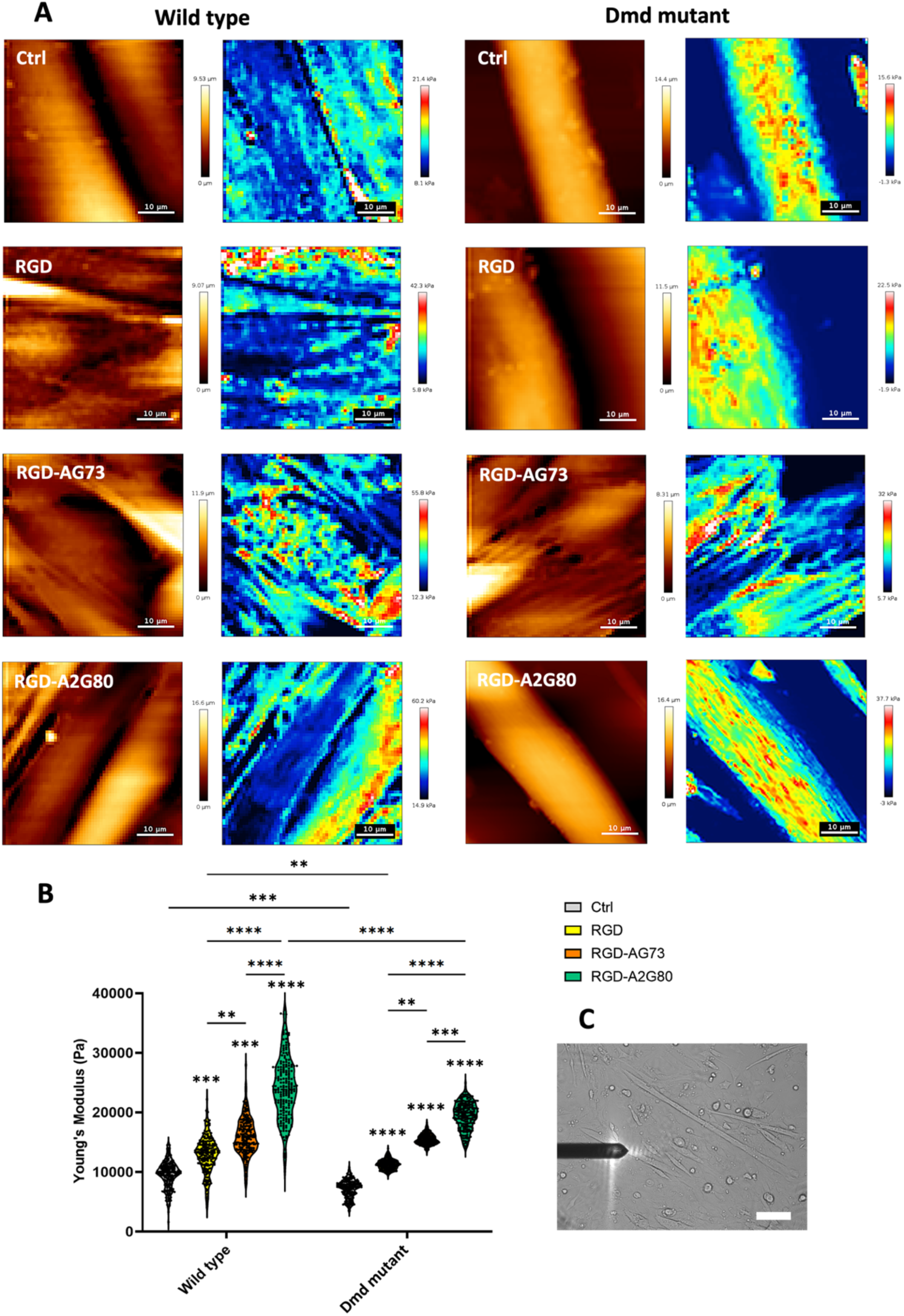
Evaluation of myotube structure and mechanical stiffness at day 7 using live Q-AFM. A) topography and corresponding stiffness map of the myotubes along with B) quantification of Young’s modulus of the myotubes and C) microscopic bright field live view of the AFM cantilever on the surface of the myotubes (scale bar: 100 µm). N=5 different positions per sample with 4000 individual stiffness data points. Asterisks represent the statistical significance between the treatment groups, with ✻ directly above data points indicating statistical differences with the control, and above the bars indicating statistical differences between the treatment groups.

The effect of peptide surfaces on myotube structure was investigated further using confocal and super-resolution microscopy (**Figures 7 and S8**). Confocal images revealed a well-defined actin cytoskeleton in both control and dystrophin-deficient myotubes, with no difference observed in actin filament length or width (**Figure 7A, B**) between genotypes. Culturing on the peptide clusters, however, increased the length and width of actin filaments in both control and dystrophin-deficient myotubes, particularly in the heterocluster samples. Accordingly, RGD-A2G80 significantly enhanced the actin filaments’ width in both cell conditions compared to all other groups. It is worth noting that although RGD-AG73 noticeably improved the actin filament length in control cells, the effect was comparable to RGD-A2G80 in the case of the dystrophin-deficient group.

**Figure 7.**
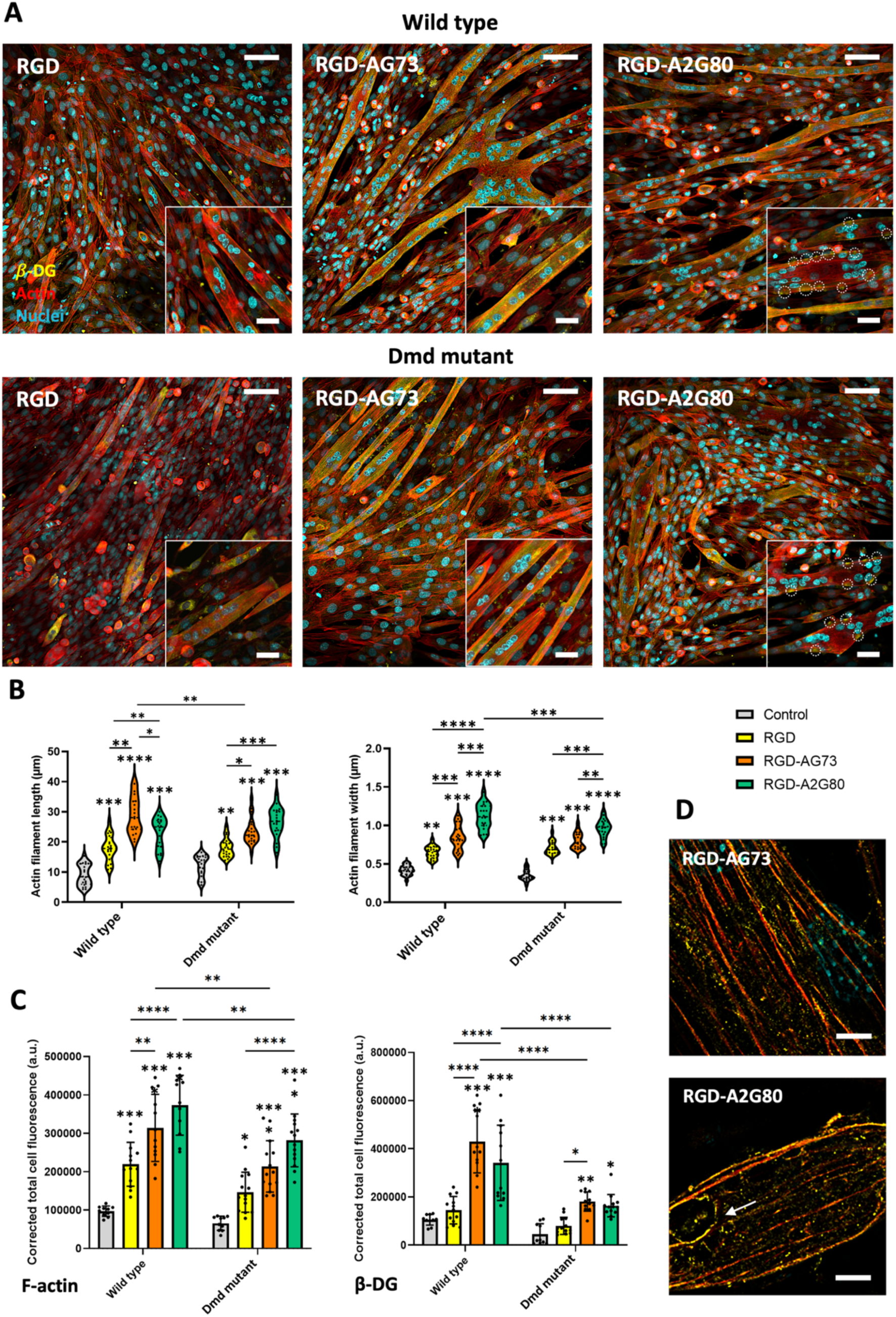
Myotube actin cytoskeletal structure and β-dystroglycan (β-DG) receptor organization analysis after 7 days. A) confocal microscopy images of the stained myotubes against F-actin (red), β-DG (yellow), and nuclei (cyan), and associated B) image analysis using the FilamentJ plugin in ImageJ software was employed for actin structural assessment, and C) further quantification of fluorescent signal related to F-actin and β-DG is depicted in graphs (n=10). D) Super-resolution microscopy images of the stained myotubes visualizing the receptor clustering and actin filaments. Scale bars in panel A are 100 and 30 µm for images and their zoomed-in version, respectively, and in panel D is 5 µm. Asterisks represent the statistical significance between the treatment groups, with ✻ directly above data points indicating statistical differences with the control, and above the bars indicating statistical differences between the treatment groups.

Similarly, the measurement of the total actin fluorescence signal revealed an increase in values associated with heterocluster compared to the control and RGD clusters in both control and dystrophin-deficient myotubes (**Figure 7C**). This is in line with the Q-AFM mechanical stiffness results, since the possible increased actin polymerization and its well-organization can improve the cytoskeletal tension along the sarcolemma of the myotubes, even with dystrophin deficiency.

Interestingly, while the total fluorescence signal for actin was comparable between control and dystrophin-deficient myotubes cultured on control surfaces, and increased with culture on the peptide surfaces, the total fluorescence signal for β-dystroglycan increased in control myotubes cultured on the RGD-AG73 and RGD-A2G80 peptide surfaces but was blunted in dystrophin-deficient myotubes (**Figure 7C**).

Closer inspection of the confocal and super resolution images suggested that the majority of the β-dystroglycan fluorescent signal was concentrated on the edge of the myotube membrane (sarcolemma) with high co-localization with actin in heterocluster surfaces (**Figure 7A)**. Surprisingly, on the RGD-A2G80 surface, β-dystroglycan signals appeared even more concentrated and formed circular clusters (unlike RGD-AG73), which have been further observed in super-resolution microscopy images (indicated by white arrows in **Figure 7A and D**) in both control and dystrophin-deficient myotubes. To the best of our knowledge, this observation has not been reported before in the literature.

As mentioned, in skeletal muscle, mechanosensitive adhesion complexes, including DAPC and focal adhesions complex (FAC), contribute to cell membrane stability by mechanically linking the actin cytoskeleton to the ECM. It has been observed that deficiencies in dystrophin components of the DAPC could negatively affect the expression and phosphorylation of focal adhesion proteins, actin polymerization, and striation ^[40]^. On the other hand, decreased FA tension has been reported in dystrophin-deficient C2C12 myoblasts due to the lack of crosstalk between dystrophin and FAs ^[35,45]^.

Our findings suggest that effective engagement with integrin, syndecan, and dystroglycan receptors is essential for structural organization and mechanical stabilization of myotubes likely via supporting DAPC-FA-mediated mechanosensation even in dystrophin-deficient condition. This study presented fundamental novel insights into the role of mechanotransduction receptors in maintaining skeletal muscle tissue homeostasis which can offer a potential target for developing muscular dystrophy treatments.

### 2.5. Culture on heterogenous peptide clusters enhanced innervation of dystrophic myotubes: synergistic effect on AChR clustering and neuromuscular junction formation

One of the implications arises from dystrophin-deficiency in skeletal muscle tissue is loss of neuromuscular connectivity and electrophysiological function due to the impaired acetylcholine receptor (AChR) clustering and postsynaptic fragmentation ^[8]^. To evaluate the effect of culturing on heterocluster peptides on neuromuscular junction (NMJ) formation in control and dystrophin-deficient myotubes we employed a previously reported sequential direct co-culturing with motor neurons for 14 days. Then, immunostaining with fluorescent α-bungarotoxin (α-BTX) which labels AChRs on myotube membranes was used to analyze their organization and clustering in both *Dmd* mutant and wild type groups (**Figure 8A**).

**Figure 8.**
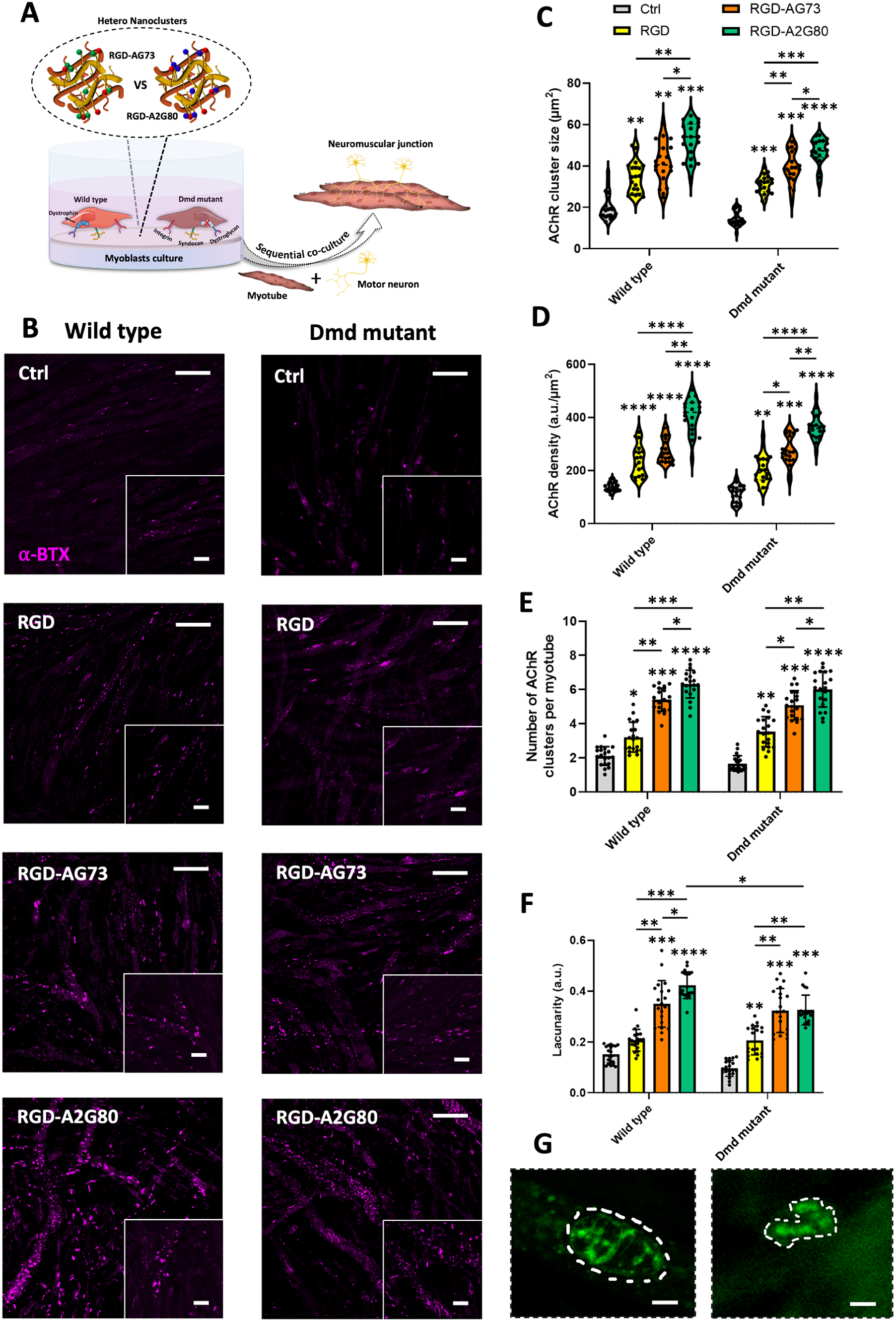
A) Schematic illustration of coculturing with motor neurons to evaluate NMJ formation *in vitro* after 14 days. B) representative confocal microscopy images of myotube stained with ⍺-BTX (magenta) which shows AChR clustering in response to peptide cluster surfaces. Image analysis using imageJ with Fraclac plugin was employed to measure AChR clusters size (C), density (D), number (E), and lacunarity (F). G) High-resolution images of AChR cluster shape in myotubes stained with ⍺-BTX (green) on RGD-A2G80 (left) and RGD-AG73 (right) cluster surfaces (cluster area is indicated with white dash lines, n=15). (scale bars are 100 and 15 µm for images and their zoomed-in version in panel B and 6 µm in panel G, respectively). Asterisks Represent the statistical significance between the treatment groups, with ✻ directly above data points indicating statistical differences with the control, unless above the bars which indicate statistical differences between the treatment groups.

As shown in confocal images and their respective quantifications (**Figure 8B-E**), in all groups AChR cluster formation was detected in both control and dystrophin-deficient myotubes cultured on all surfaces. However, our engineered biomaterial surfaces with heterogenous ligand clustering significantly enhanced the number and size of AChR clusters per myotube, with increased associated fluorescent intensity signals per area of cluster (AChR density) compared to control. Specifically, the heterogenous peptide cluster surfaces produced approximately 2.5-3 times more clusters than control and 1.7 times more than RGD homogenous cluster. Furthermore, the largest average cluster size was observed in RGD-A2G80 samples with ∼52.4 µm^2^.

Previous studies by us and others have shown the importance of AChR cluster shape and formation of lacunas as one of the hallmarks of postsynaptic organization during NMJ development ^[46–48]^. Therefore, we analyzed the spatial heterogeneity of AChR clusters using fractal analysis of confocal images and measurement of lacunarity index (**Figure 8F**).

Accordingly, in both *Dmd* mutant and wild type groups, AChR clusters formed on the control surface were small (≤20µm^2^) with low lacunarity, suggesting early-stage receptor clustering and limited NMJ development. In contrast, in addition to increased cluster density, number, and size, peptide cluster surfaces noticeably improved lacunarity, particularly in heterogenous cluster surfaces than those on the control. Specifically, in RGD-A2G80 group, AChR clusters displayed a perforated and almost pretzel shape (**Figure 8G**) structure with the highest lacunarity (**Figure 8F**) compared to RGD-AG73 group with fewer lacunas and a c-shaped shape. Such receptor arrangement can be associated with later stages of postsynaptic machinery development and NMJ formation, with neurotransmission and synaptic function likely due to the enhanced β-dystroglycan-AChR mediated transmission of ACh and agrin, as the main neurotransmitters ^[8,48]^ (although this remains to be investigated), along with improved sarcomere organization and myotube development.

We then conducted confocal microscopy and immunostaining against markers for neuronal microtubules (β-Tubulin III, yellow), myotube myosin heavy chain (MF20, magenta), and AChRs (α-BTX, green) to visualize the neuromuscular connections and further quantified the neurogenesis and number of junctions (**Figures 9 and S9**). According to **Figure 9A**, in *Dmd* mutant myotubes on the control surface, there is limited neurite formation and connectivity with myotubes, with colocalization Pearson’s correlation coefficient of ∼0.05. However, without the need for neurotrophic factors, culturing on heterogenous peptide cluster surfaces significantly improved neurogenesis and neuromuscular connection with colocalization Pearson’s correlation coefficient of ∼0.82 compared to control and RGD cluster surface in both wild type and *Dmd* mutant groups (**Figures 9A and S9**).

**Figure 9.**
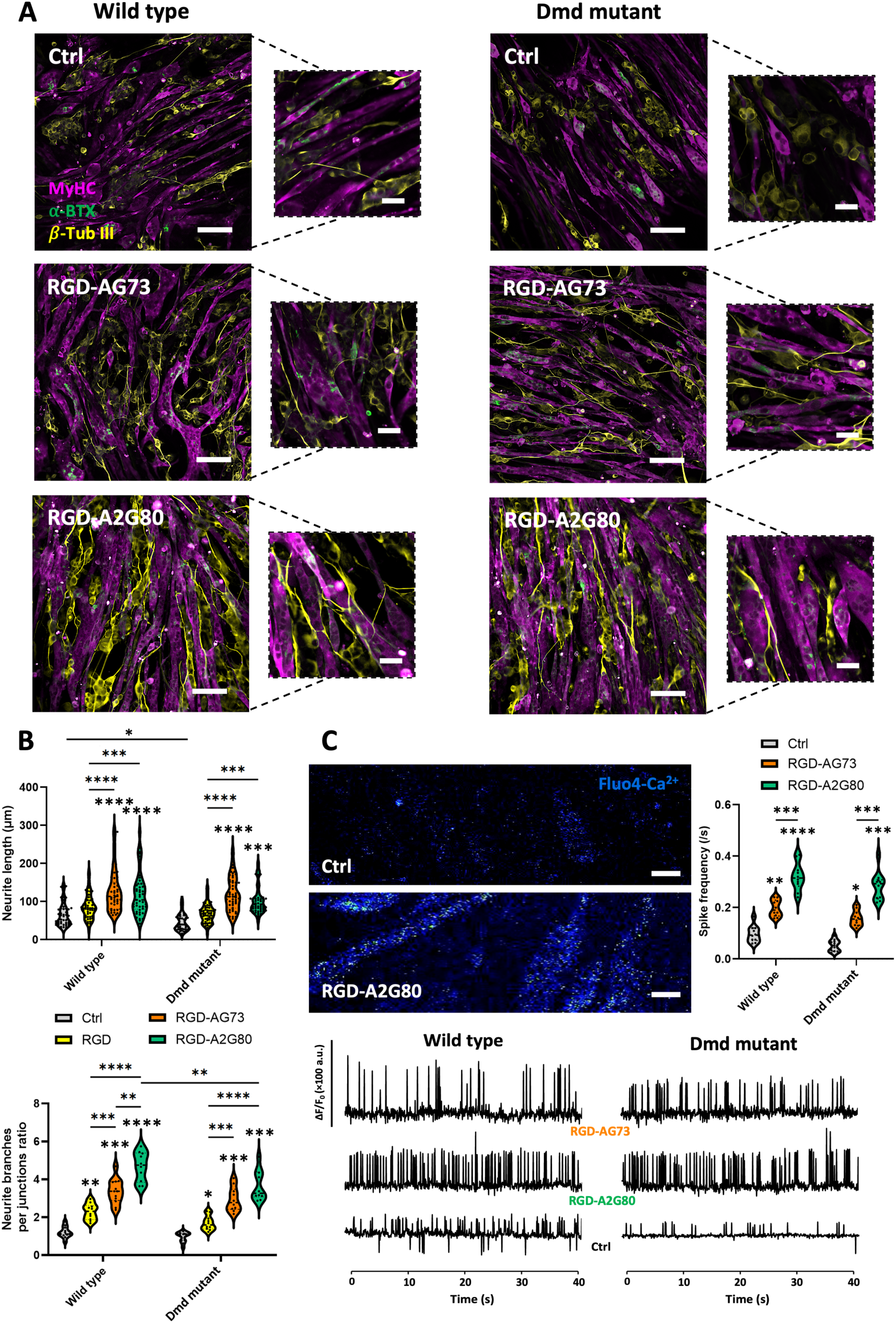
Evaluation of neuromuscular connectivity after 14 days of culture on peptide-clustered surfaces in response to the ligand type. Representative A) confocal images of neuromuscular junction stained against MyHC (MF20, magenta), AChR (⍺-BTX, green), and β-tubulin III (yellow), scale bars: 100 and 30 µm for zoom-in panel. B) Image analysis using NeuronJ plugin of ImageJ software was performed to measure neurite length and ratio of neurite branches per junction. C) Confocal microscopy imaging (resonant mode with 20 fps) for calcium transient analysis in myotubes stained with the Ca²⁺ indicator Fluo-4 AM under the effect of ligand type in heterocluster surfaces. Ca transition recorded for 1-2 min, and then the signal intensity of images and spike frequency were quantified using ImageJ Z-profile and Time series analyzer plugin, respectively (scale bars: 30 µm and n=20, 15, and 10). Asterisks represent the statistical significance between the treatment groups, with ✻ directly above data points indicating statistical differences with the control, unless above the bars which indicate statistical differences between the treatment groups.

Although there was no noticeable difference in neurite length between RGD-AG73 and RGD-A2G80, the ratio of neurite branches per junctions, which represents the quality of myotube innervation, was slightly higher in RGD-A2G80 suggesting increased synapse formation (**Figure 9B**). Interestingly, the neurite organized almost in parallel to the direction of myotubes under the effect of peptide cluster surfaces and this alignment was further seen in RGD-A2G80. It is hypothesized that such efficient axonal guidance and synaptic targeting can be the result of regulating cytoskeletal organization, which facilitates the subsequent contractile function of the *in vitro* formed muscle construct.

These results can be explained by the decisive role of DAPC and FA in maintaining structural integrity of the myotubes and stabilization of synaptic machinery. In this regard, previous studies have shown that effective engagement of integrins (particularly α_7_β_1_, α_3_β_1_, and α_5_β_1_) and FA-mediated signalling plays a key role in AChR clustering during NMJ formation by not only regulating the membrane structure of myotubes but also supporting synaptic integrity for assembling the postsynaptic ^[49–52]^. For example, it has been shown that involvement of integrin α_5_β_1_ in signalling of DAPC via FAK or Src protein secretion can also increase synapse stabilization and postsynaptic differentiation ^[4]^.

Moreover, dystroglycan close interaction with AChR ^[4]^ and their effect on promoting cytoskeletal alterations, synapse function, clustering of AChRs and functional maintenance ^[5,6]^ has been suggested, which are essential for the development of neuromuscular junctions ^[8]^. On the other hand, neural patterning and axonal guidance are also regulated by multiple signalling molecules engaged in integrin-syndecan interactions; for example, altering PKC and FAK activity has an impact on neurite formation and axonal branching ^[8,10]^.

As a measure of maturity and functionality in co-culture condition, myotube contractility was assessed via calcium imaging using Fluo4-AM as a fluorescent calcium indicator (**Figure 9C and Supplementary Information Movies**). In both control and dystrophin-deficient myotubes, culture on the heterocluster surfaces improved the generation of regular calcium transient spikes in response to ACh administration, with sharp and continuous peaks and high amplitude, compared to the control condition with low amplitude calcium transients (**Figure 9C**). Interestingly, the calcium transient pattern was more regular and synchronized with almost the same amplitudes, particularly in RGD-A2G80 with an increased proportion of spontaneous beating in both mutant and wild type groups. Further analysis of the calcium spike frequency in the myotubes formed on the different surfaces demonstrated a significant increase in RGD-A2G80 (0.31 peak/s) compared to RGD-AG73 (0.18 peak/s) and control (0.07 peak/s) groups (**Figure 9C**), which may be related to the regulating role of syndecan-4. Accordingly, despite an increase in expression of syndecan-4 receptors in the myoblasts promoting proliferation, in later phases of myotube development, these receptors underwent internalization ^[3,53]^. We hypothesize that this event may weaken the signalling effect of binding to AG73 peptides within the heterocluster during myotube maturation, affecting contractility. These findings highlight the effect of peptide clusters and associated receptor crosstalk on increasing myotube maturity and resulting in robust contractility even without external electrical stimulation.

Previous studies indicated the crucial role of calcium transients in regulation of myogenic proteins for skeletal muscle development, and disrupted calcium transients in diseased muscle can impair myogenesis, myotube organization, and sarcomere assembly ^[54–56]^. In line with these previous studies, we observed enhanced fusion index and improved sarcomere and cytoskeleton organization in myotubes formed on heterocluster surfaces and improved synchronized calcium transient peaks suggesting enhanced myotube formation even in dystrophin-deficient cells.

We have shown that stabilizing and organizing DAPC with increased formation of FA via facilitated crosstalk between the main involved receptors offers a promising and unexplored strategy for promoting dystrophic skeletal muscle tissue regeneration *in vitro*. Our study is among the first to exploit the synergistic effect of integrin-syndecan and integrin-dystroglycan signalling using engineering receptor binding ligands presentation on biomaterial surface to improve the innervation of skeletal muscle even in the absence of dystrophin. However, further studies, including proteomic analysis are required to uncover the changes in the marker protein secretion as a result of culturing on this engineered biointerface. In addition, employing controlled electrical stimulation bioreactors can be considered for achieving a higher level of tissue maturation required for diseased tissue modelling.

## 3. Conclusion

Dystrophin deficiency in skeletal muscle tissue negatively impacts myogenesis and structural integrity of the myotubes, which affects their functionality. In the current study, we investigated the impact of increased DAPC and FA formation and their targeting receptor crosstalk on the improvement of skeletal muscle tissue regeneration in dystrophic conditions. This was achieved by engineering biointerface of skeletal muscle cells via functionalization of a RAFT synthesized low-fouling copolymer with extracellular matrix protein-derived adhesion ligands and their clustering at the biomaterial biointerface. The results confirmed the synergistic effect of integrin-syndecan/dystroglycan engagement and their clustering on ameliorating the lack of myoblasts’ adhesion, proliferation, and myogenesis caused by dystrophin deficiency. Moreover, our results indicated the crucial role of improved focal adhesion and increased receptor localization, particularly dystroglycan, at the sarcolemma, on enhancing myotube structural organization, mechanical integrity, and neuromuscular connection, which are necessary for proper muscle functions. This can be a promising approach for tissue engineering and regenerative therapies, where biomaterials functionalized with such peptides could promote muscle repair and functional recovery in disease contexts or post-injury and build personalized disease tissue models for drug screening studies.

## 4. Experimental Section

Materials and methods associated with the polymer synthesis and characterization are presented in the **Supplementary Information**.

### 4.1. Sample Preparation

Based on our previously published optimizations ^[14]^, blends of functionalized polymer (MPP, with local densities of ∼4 peptides per chain) and non-functionalized polymer (MP) were prepared using appropriate blending ratios to reach global densities of 7 µg of peptide per mg of polymer mixture and dissolving the polymers with 10 wt.% in methanol:water (4:1) at 60 °C. Non-functionalized polymer (MP), randomly distributed RGD (rRGD), and polymer blend with scrambled peptides (ScPEP, containing RGE-A2G80SC-AG73SC) surfaces were employed as the control surfaces. To generate polymer film, polymer solution was cast (50 µl.mm^-2^) onto substrates (round glass coverslips, No.1, 11 mm) and then samples were slowly dried under a fume hood at room temperature for 24 h before use. For biological assays, all samples were sterilized using UV light for 2 h, 3× rinsing with sterile PBS, and then incubated with serum-free culture media at 37°C for overnight before cell seeding. The details of surfaces are summarized in **Table 2**.

### 4.2. Raman Spectroscopy

To confirm the presence and covalent conjugation of peptides at the interface, FTIR-ATR spectroscopy was performed using a Bruker LUMOS Fourier-Transform Infrared Spectrometer (Bruker Corp). For each surface, a measurement involving 32 scans at a resolution of 4 cm^−1^ was performed.

### 4.3. Energy Dispersive X-ray Spectroscopy (EDX)

EDX microanalysis was used to measure the amount of nitrogen at the surface of polymer films, indicative of the presence of nitrogen arising from the peptide bonds within the ligands. EDX analysis was performed using an energy dispersive spectroscopy module (Bruker) coupled to a scanning electron microscope (Hitachi FlexSEM 1000, Japan) using an accelerating voltage of 20 keV. Spin-cast polymer films were exposed to DI water overnight and then lyophilized at −20°C prior to EDX analysis.

### 4.4. Kelvin Probe Atomic Force Microscopy (KPFM)

Atomic force microscopy in KPFM mode (Asylum Research Cypher, USA) were employed for measurement of surface charge of spin coated peptide-functionalized polymer films on silicon wafers after reaching equilibrium swelling in MilliQ water. Probing the surface was conducted with a platinum-iridium coated cantilever (FMG01/Pt, TipsNano) with an apex radius of approximately 35 nm and a resonance frequency of 71.7 kHz. The samples were blown with nitrogen stream before test in air and they were electrically grounded by applying conductive silver ink to ensure a stable reference during measurements. Topography (semi-contact mode) and surface potential (tapping mode) maps were simultaneously captured over a scan area of 5 × 5 µm^2^ and analyzed based on the electrical voltage applied to disable the mechanical excitation of cantilever’s resonance. The data collected from 5 different points of the surfaces with 3 replicates.

### 4.5. Cell Culture and Maintenance

Murine myoblasts (C2C12s, ATCC, Manassas, VA, USA) were genetically modified with the *CRISPR/Cas9* single guide RNA, knocking down the dystrophin protein expression to generate *DMD* C2C12 (*Dmd* mutant) cells or without the target protein to generate control (wild type) cells as described in the Supplementary Information. Dystrophin knockdown was validated in myotubes via western blot and immunostaining against dystrophin (**Figure S6**). Non-modified C2C12 myoblasts were also employed for the preliminary evaluation of focal adhesion response, illustrated in the Supplementary Information. Cells were grown in growth media (GM) containing Dulbecco’s Modified Eagle Medium-High Glucose (DMEM-Glutamax, Gibco, Invitrogen) supplemented with 10% v/v Fetal Bovine Serum (FBS, Gibco, Invitrogen) and 100 units.ml^-1^ penicillin and streptomycin (P/S, Gibco, Invitrogen) and incubated at 37 °C in a humidified, 5% CO_2_ atmosphere. The culture medium was changed every 2 days, and cells at 70% confluency (passage numbers 7-10) were used for seeding.

Murine motor neurons (NSC-34, ATCC, Manassas, VA, USA) were maintained in GM (DMEM-Glutamax supplemented with 10% v/v FBS, and 100 units.ml^-1^ P/S) and incubated at 37 °C and 5% CO_2_ atmosphere. The culture medium was changed every 3 days, and cells at 90% confluency (passage numbers 8-10) were used for seeding. For neuronal differentiation, after 2 days, the growth media were changed to DM (Neurobasal medium (NBM) supplemented with 2% v/v B-27, 0.25% v/v L-glutamine, 1 μM retinoic acid, and 100 units.ml^-1^ P/S) and differentiated for 5 days.

### 4.6. Myoblast Cell Morphology and Proliferation

Myoblasts or motor neurons were seeded on samples in ultra-low attachment 24-well plates (Corning) with 2×10^4^ cells per well. After 24 h, cell adhesion and morphology were evaluated by immunostaining. Briefly, PBS-washed samples were fixed in 4% paraformaldehyde for 20 minutes, washed 3 times with PBST (PBS+ 0.3% Triton X-100 (Sigma)) and blocked for 1 h in 4% BSA solution in PBST. Then, samples were incubated in a solution of rabbit polyclonal anti-vinculin (1:250, PA5-29688, Invitrogen) diluted in PBST and 2% BSA at 4 °C overnight. After 3×10 min PBS wash, samples were incubated in a solution of anti-rabbit Alexa Fluor 488 (1:500, A21206, Invitrogen) for 2 h, followed by 1h incubation in ActinRed 555 Ready Probes (R37112), and 10 min in DAPI (1:1000, D3571) (Life technologies) and final wash with PBS for 4 times. Samples were imaged using a Nikon A1R+ confocal laser scanning microscope (Japan, 20x magnification). The cell spreading area and orientation were analyzed with ImageJ ^[57,58]^.

The alamarBlue assay was employed to assess cell proliferation for 3 days. Briefly, 1×10^4^ cells per well of 24-well plates were seeded on samples and each day, GM was removed and replaced by fresh GM containing 10% v/v solution of 140 µg.ml^-1^ resazurin in PBS (Sigma) and incubated for 3 h. Fluorescence intensity of each well was recorded using TECAN infinite M200PRO microplate reader (Ex/Em wavelengths: 530/590 nm). The fluorescence intensity was then converted to cell density using a standard equation generated from absorbance data of multiple cell densities.

### 4.7. Myoblast Differentiation into Myotubes

The myoblasts were seeded on samples at a density of 1.5×10^4^ cells.cm^-2^. After 3 days in GM, the media changed to differentiation medium (DM) containing DMEM-Glutamax supplemented with 2% horse serum (HS, Gipco, Invitrogen) and 1% P/S, and maintained for differentiation to myotubes for 7 days (media changes every 2 days). At days 7 of differentiation, cells were stained following the previously described procedure using overnight incubation in solutions of MYH1E antibody (MF 20)-DSHB (uiowa.edu), α-Actinin primary antibody (EA-53, MFCD00164521, Sigma), and dystrophin MANEX1011B(1C7)-DSHB (uiowa.edu) at 1:100, 1:250, and 1:100 dilutions, respectively and subsequent incubation in secondary antibodies solution of anti-mouse Alexa Fluor 555 (1:500, A-21422, Invitrogen) and anti-mouse Alexa Fluor 488 (1:500, A-11001, Invitrogen) and DAPI (1:1000) for 2 h before imaging with the Nikon A1R+ confocal laser scanning microscope (20x magnifications). ImageJ analysis based on the method mentioned in ^[59,60]^ was employed to measure fusion index, myotube diameters, orientation, aspect ratio, and the ratio of sarcomere-forming myotubes. Analysis of sarcomere length and uniformity was conducted using SotaTool software.

### 4.8. Myotube Stiffness Measurement Using Q-AFM Nanoindentation

The Nanowizard IV (JPK Instruments, Germany) Q-AFM was used to assess cell structure and dynamics. Measurements were taken in L15 medium at 37°C in a PetriDishHeater (Bruker, Germany), using a ContGD-G cantilever (BudgetSensors, Bulgaria) with a spring constant of 0.2 N.m^-1^. JPK software was used to fit all force-distance curves with the Hertz model using a parabolic indenter, with a fitting range of less than 18% of cell thickness, to produce the myotube Young’s modulus. Contact points were calculated with the zero-gradient approach. Young’s modulus map was generated by analyzing more than 3000 individual force-distance curves over a scan area of 50 × 50 µm^2^. The reported data were collected from at least 5 different points in the samples.

### 4.9. Analysis of Actin Cytoskeleton and Receptor Localization

Then cells were stained following the previously described procedure using overnight incubation in solutions of β-dystroglycan primary antibody (clone 7D11, MABT1546, Sigma) and at 1:250 dilution, and subsequent incubation in secondary antibody solution of anti-mouse Alexa Fluor 647 (1:500, ab150115, Abcam) and Alexa Fluor 555 conjugated actin phalloidin (A21206, Invitrogen) for and DAPI (1:1000) for 2 h before imaging with the Nikon A1R+ confocal laser scanning microscope (20x magnifications). ImageJ was used for fluorescent signal measurement and FilamnetJ was employed to analyze actin.

The super-resolution microscope Elyra 7 (Zeiss with Lattice SIM^2^) with 60x magnification was also employed to visualize the clustering of the β-Dystroglycan within the myotubes membrane on the heterocluster surfaces.

### 4.10. NMJ Formation and Evaluation

Myotube and motor neuron co-culture were carried out based on our previous method. Accordingly, differentiated neurons were gently trypsinized (1:0.5 ratio PBS diluted trypsin, Gipco, Invitrogen) and seeded on top of the grown myotubes (day 7) on the samples with a density of 1 × 10^4^ cells/cm^2^ with 1:1 mixture of neuronal and myotube DMs. After one day, the co-culture media changed to neuronal DM supplemented with 1% v/v HS until the end of the experiment (Day 14).

Staining with α-bungarotoxin was used to visualize and quantify the average number, size, and density of AChR clusters in co-culture samples (based on ^[46]^). To label AChRs, the previously mentioned immunostaining process was applied using α-bungarotoxin-Alexa Fluor 488 (1:500, Invitrogen) in a secondary antibody solution and incubated for 2 h. The neurites of differentiated neurons were also immunostained with an antibody solution of mouse monoclonal anti-beta tubulin III (1:2000, G7121, Promega) and subsequent staining with anti-mouse Alexa Fluor 647 (1:500, A-21235, Invitrogen). The colocalization of immunolabeled myotubes/neurons and AChR was conducted by JACoP plugin of ImageJ in which the Pearson’s correlation coefficient higher than 0.5 was considered as perfectly overlapped fluorescent signals.

Images were captured using the Nikon A1R+ confocal microscope and at 20x, 60x, and 100x (oil immersion lens) magnifications, from at least five random positions on each sample. The ImageJ particle analyzer was used to identify AChR cluster outlines (by binarizing the cluster outlines), and clusters less than 4.5 μm² were removed from further analysis. Cluster numbers were calculated by counting AChR clusters in each image close to neurites and normalising them to the number of MyHC-positive myotubes. The size and density of AChR clusters were measured and averaged across groups using ImageJ regions of interest (ROI) and fluorescent intensity analysis. Additionally, to evaluate morphological heterogeneity between AChR clusters on different surfaces, fractal analysis of α-BTX-stained samples was performed using the FracLac plugin in ImageJ ^[46]^. Accordingly, the AChR clusters outline were binarized, and lacunarity was calculated and averaged for each experiment.

### 4.11. Calcium Transient Imaging

At day 14, cells were labelled with cell-permeable calcium indicator Fluo-4 AM (F14201, Invitrogen) following the manufacturer’s protocol. Briefly, the cells were incubated at 37 °C with 1 µM of Fluo-4 AM dye solution mixed with 1 μl of 20% Pluronic F-127 solution in DMSO in neurobasal media without phenol red. After 1 h, they were rinsed twice with HBSS and allowed to recover in the incubator for 15 min before imaging. Samples were then imaged in a glass-bottom opaque 24-well culture plate while incubating in HBSS at 35±3 °C. Fluorescent time-series data were collected using Nikon A1R+ confocal microscope (20x magnification) in resonant mode, capturing calcium transients for 1-2 min at 20 fps after treatment with 10 µM acetylcholine (Sigma Aldrich). ROIs were identified manually in areas displaying calcium signals, and fluorescent intensity in different time frames was analyzed using ImageJ (Z-axis profile). During analysis, the intensity values were subtracted from the background intensity, and the averaged fluorescent intensity was plotted against time. Additionally, after applying the Gaussian filter, the calcium spike frequency was quantified using the Time Series Analyzer plugin and Find Peaks macro.

### 4.12. Statistical Analysis

Data in this study are reported as mean ± standard deviation and using GraphPad Prism 9 software, they were graphed and statistically analyzed through one-way and two-way ANOVA via the Tukey-Kramer multiple comparison test (significance level of p-value < 0.05). Unless otherwise indicated, all tests were carried out with at least three independent experiments and technical/biological repeats (n). Statistical significance is indicated as p ≤ 0.05 (*), ≤ 0.01 (**), ≤ 0.001 (***), and ≤ 0.0001 (****).

## Supplementary Information

Supplementary Information is available from the author or the Wiley Online Library.

## Acknowledgements

S.N. gratefully acknowledges the support of the University of Melbourne via the Melbourne Graduate Research Scholarship. The authors would like to appreciate the Protein Characterization and Peptide Synthesis Platform at Bio21 Institute and the Materials Characterization and Fabrication Platform (MCFP) at the University of Melbourne, particularly Dr. Paul Brannon, for assistance with the image acquisition from the super-resolution microscope.

## Data Availability

Data will be made available on request from authors.

