## Supporting Information for "Changing the Fate of Dystrophin-Deficient Myoblasts via Hetero Ligand Nanoclusters on Biomaterial Surface: Effects of Integrin-Syndecan or Dystroglycan Crosstalk"

### 1. Experimental Section

#### 1.1. Materials

Methyl methacrylate (99%), Poly(ethylene glycol) methyl ether methacrylate (PEGMA, Mw: 475 g.mol-1), PEGMA-OH ($M_w$ = 668 g.mol$^{-1}$) synthesized monomer, 2,2′-Azobis(2-methylpropionitrile) (AIBN), triethylamine (TEA), 4-Dimethylaminopyridine (DMAP), 5-Norbornene-2-carboxylic acid (mixture of endo and exo, predominantly endo), N-(3-Dimethylaminopropyl)-N′-ethylcarbodiimide hydrochloride (EDC-HCl), 2-phenyl-2-propyl benzodithioate (99% RAFT agent), Azobisisobutyronitrile (AIBN, RAFT initiator), tris(2-carboxyethyl) phosphine, 2,2-dimethoxy-2-phenylace Mimotopes Pty Ltd (Clayton, Victoria, Australia) supplied the amino acids, resin, N,N′-diisopropylcarbodiimide (DIC), and Oxyma used in peptide synthesis. Unless otherwise specified, these compounds were utilised without any purifying procedures. To remove inhibitors, the MMA and PEGMA were passed through basic alumina columns and kept at temperatures ranging from 2 to 8 °C. The hydroxyl-terminated poly (ethylene glycol) methacrylate (PEGMA-OH, Mw = 668 g/mol) monomer was synthesised using the method described in our previous work [1,2].

#### 1.2. Methods

##### *1.2.1. Peptide synthesis*

The adhesive peptides have been synthesized using Fmoc-based solid-phase peptide synthesis method for amide C-terminated peptides on Rink amide resin (resin loading: 0.649) and CEM Liberty Blue microwave peptide synthesiser based on the protocol in [1]. For the conjugation of peptides to polymer via thiol-ene click chemistry, cysteine amino acid was incorporated into the N-terminus sequence of peptides along with additional glycine amino acids providing physicochemical freedom for peptide sequences in contact with cells (**Table 1 and Figure S1**).

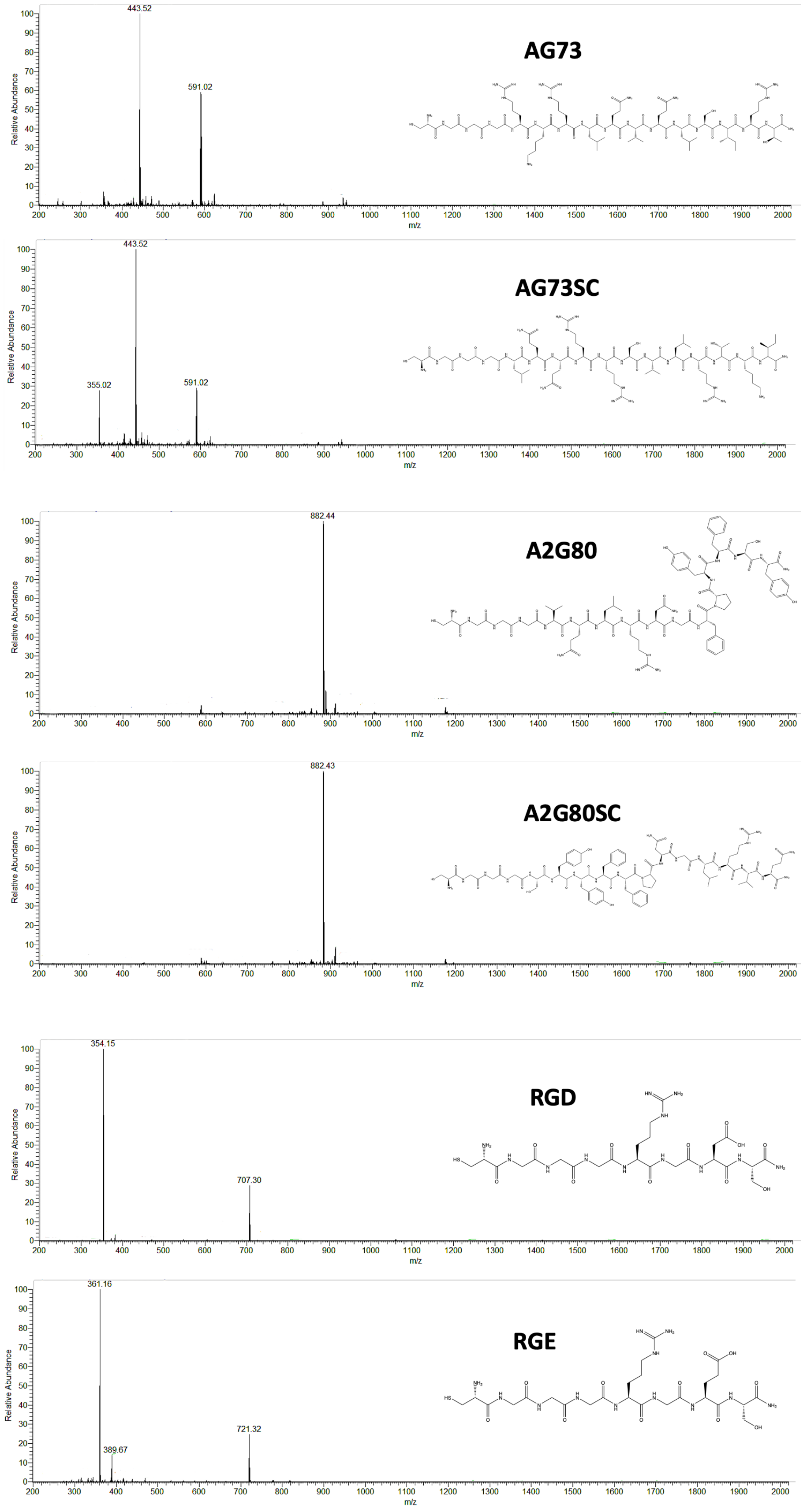

**Figure S1.** Mass spectrum of the purified AG73 (Exact Mass: 1770.02, Observed Mass: 591.02 (M+3H), 443.52 (M+4H)), AG73SC (Exact Mass: 1770.02, Observed Mass: 591.02 (M+3H), 443.52 (M+4H)), A2G80 (Exact Mass: 1762.82, Observed Mass: 882.44 (M+H)), A2G80SC (Exact Mass: 1762.82, Observed Mass: 882.43 (M+H)), RGD (Exact Mass: 706.28, Observed Mass: 707.30 (M+H), 354.15 (M+2H)), RGE (Exact Mass: 720.30, Observed Mass: 721.32 (M+H), 361.16 (M+2H)).

#### *1.2.2. Polymer synthesis and sample preparation*

The synthesis and functionalization of polymer have been done based on the previously mentioned protocol using RAFT polymerization and thiol-ene click chemistry [1]. The evaluation of polymer has been carried out using GPC in DMF and $^1$H NMR (**Figure S2**). To form heteroclusters, during thiol-ene click reaction, 50:50 mol% ratio of RGD:AG73 or RGD:A2G80 were used.

To determine the composition of the monomer and polymers in deuterated chloroform, proton nuclear magnetic resonance (1H NMR) spectroscopy was performed using a Varian Unity Plus 500 MHz NMR spectrometer (CDCl3, 99.8% purity from Cambridge Isotope Laboratories). GPC analysis was performed using a Shimadzu liquid chromatography system and three Phenomenex Phenogel columns (with porosities of 500, 104, and 106 Å; bead size: 5 mm). The columns were kept at 45 ± 1°C. An interferometric refractometer, the Wyatt OPTILAB DSP (functioning at 633 nm), was used. The mobile phase was dimethylformamide (DMF) of high-performance liquid chromatography (HPLC) quality, which flowed at a rate of 1 ml/min. The number-average molecular weight (Mn), weight-average molecular weight (Mw), and polydispersity index (PDI) of the polymers were calculated by calibrating the system with tightly dispersed poly(methyl methacrylate) standards.

**Figures S2, 3** show the $^1$H NMR and GPC characteriszations of synthesised polymers in deuterated chloroform and DMF, respectively. Following the polymer purification procedure, the molar ratio of MMA to PEGMA was established by analysing their unique peaks in the $^1$H NMR spectra (see equations in **Table S1**). In MP equations, $X_{PEGMA}$ and $X_{MMA}$ represent the mole percentages of PEGMA and MMA residues in the polymer, respectively. $I_C$ is the integral area corresponding to the $CH_2$ peaks of the $COOCH_2$ methylene group protons inside the repeating unit of PEGMA, whereas $I_A$ is the integral area associated with the $CH_3$ peak for the $CH_3$ methyl group protons in the backbone of both MMA and PEGMA.

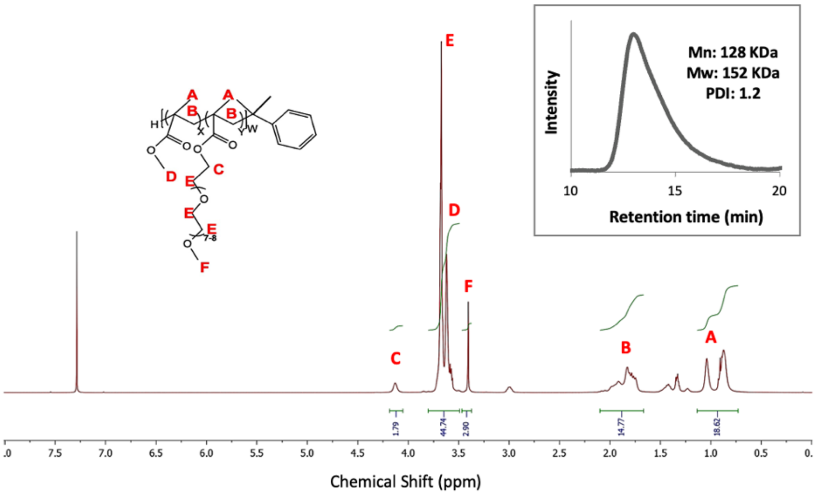


**Figure S2.** $^{1}$H NMR spectrum in deuterated chloroform of MP polymer with GPC spectrum in DMF.

Based on Fig. S1, for MP polymer, $^{1}$H NMR (CDCl$_3$) includes δ 4.15 (s, 3H), δ 3.62-3.8 (m, 3H), δ 3.5-3.62 (m, 1H), δ 3.4 (s, 1H), δ 1.6-2.1 (m, 1H), δ 0.68-1.12 (d, 1H).

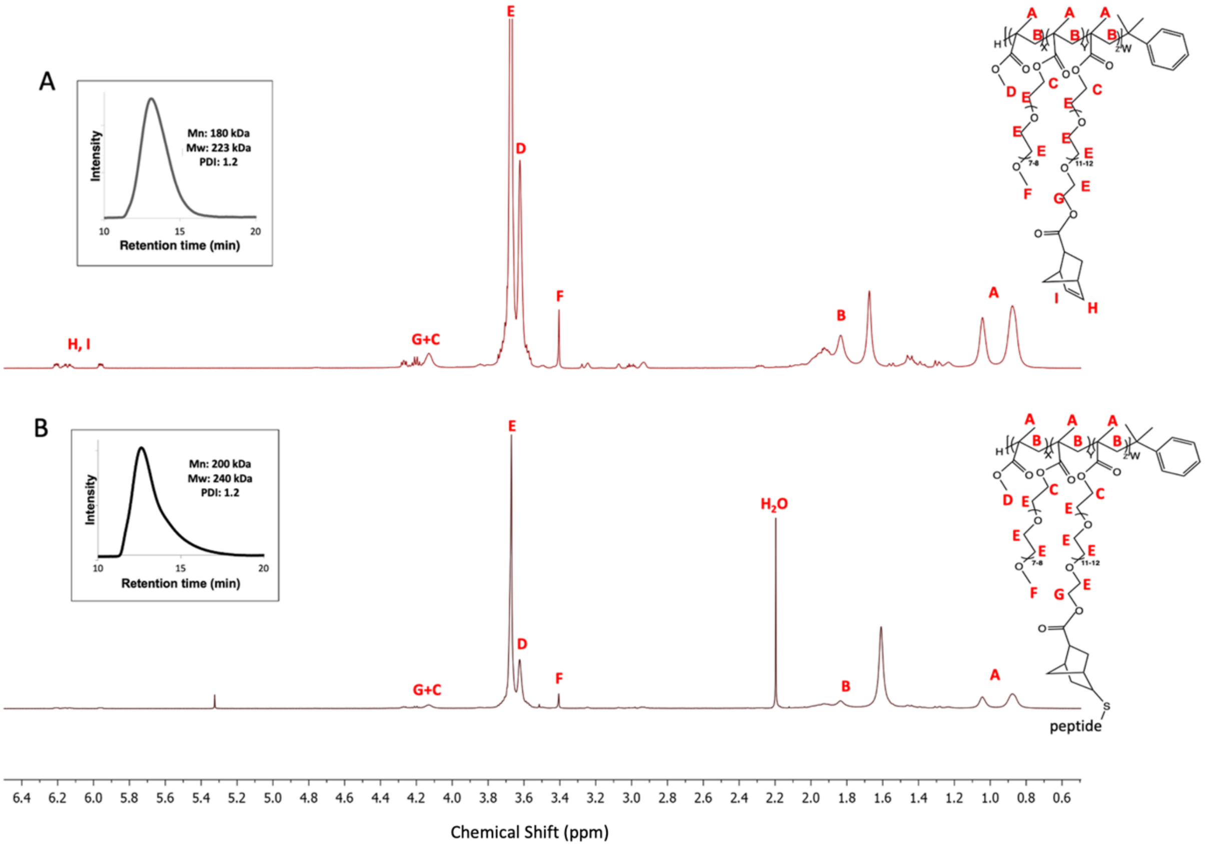


**Figure S3.** GPC spectrum in DMF and $^{1}$H NMR spectrum in deuterated chloroform of A) norbornene functionalized MPP polymer and B) after peptide conjugation. After peptide conjugation, the picks around 6.0 ppm which is related to the conjugation of norbornene disappear.

In MPP calculations, $X_{PEGMA+PEGMAOH}$ and $X_{MMA}$ represent the mole percentages of PEGMA+PEGMAOH and MMA residues in the polymer, respectively. $I_C$ is the integral area associated with the $CH_2$ peaks that correspond to the $COOCH_2$ methylene group protons inside the repeating units of PEGMA+PEGMAOH. In contrast, $I_A$ is the integral area of the $CH_3$ peak corresponding to the $CH_3$ methyl group protons in the backbone of MMA, PEGMA, and PEGMAOH. Furthermore, $X_{PEGMA}$ and $X_{PEGMAOH}$ represent the mole percentages of PEGMA and PEGMAOH residues in the polymer, respectively. $I_F$ is the integral area associated with the $CH_3$ peaks that correspond to the $CH_3$ methyl group protons inside PEGMA's repeating units.

#### 1.2.3. Evaluation of norbornene functionalization of polymers and conjugation of synthesized peptides to polymer

After norbornene conjugation $^1H$ NMR ($CDCl_3$) results include δ 6.18-6.22 (dd, 3H), δ 6.10-6.17 (m, 3H), δ 5.90-5.97 (dd, 3H), δ 4.02 (s, 3H), δ 3.52-3.75 (m, 3H), δ 3.41-3.52 (m, 1H), δ 3.3 (s, 1H), δ 1.62-1.88 (m, 1H), δ 0.65-1.2 (d, 1H).

**Table S2** shows the composition of the NB functionalized polymers, as well as the formulae used to calculate the $^1H$ NMR and GPC data. According to the formulae, $I_{PEGMA+PEGMANB}$ represents the total integral area of the PEG monomers inside the polymer. Furthermore, $I_{G+C}$ represents the integral area ascribed to the $CH_2$ peaks of the $COOCH_2$ methylene group protons in PEGMA+PEGMANB. The presence of 2/3 represents the ratio of $CH_{2,C}$ to the total number of $CH_{2,C}$ and $CH_{2,G}$ in PEGMA+PEGMANB. $I_{PEGMAOH}$ denotes the integral area connected to PEGMAOH, whereas $X_{PEGMAOH}$ and $X_{PEGMA+PEGMAOH}$ reflect the mole percentages of PEGMAOH and PEGMA+PEGMAOH in the polymer, respectively. These values are determined from the equations in Table S1. $X_{PEGMANB}$ represents the mole percentage of PEGMANB in the polymer. $I_H$ and $I_{I'+H'}$ denote the integral regions associated with norbornene's CH peaks in PEGMANB residues.

**Table S1.** Composition of polymers used in this study.

| **Polymer name** | **Time of polymerization (h)** | **Conversion** | **Polymer composition before conjugation (mol%) (MMA-** | **Polymer composition after conjugation (mol%) (MMA-PEGMA-** | **$M_n$ (kDa)** | **PDI** |
|---|---|---|---|---|---|---|

|  |  |  | PEGMA-PEGMAOH) | PEGMAOH-PEGMA-NB) |  |  |
|---|---|---|---|---|---|---|
| **MP** | 17 h | 60% | 84.4-15.6-0 | - | 128 | 1.2 |
| **MPP** | 17 h | 67% | 83.7-2.3-14.0 | 83.7-2.3-1.4-12.6 | 180 | 1.2 |

The actual polymer compositions were calculated based on H NMR data using the equations below:

**MP:**

$$X_{PEGMA} = \frac{\frac{I_C}{2}}{\frac{I_A}{3}} \times 100 \qquad X_{MMA} = 100 - X_{PEGMA}$$

**MPP:**

$$I_{PEGMA+PEGMANB} = \frac{I_{G+C}}{2} \times \frac{2}{3} \qquad X_{PEGMANB} = X_{PEGMAOH} \times \frac{I_H + \frac{I_{I'+H'}}{2}}{I_{PEGMAOH}}$$

$$I_{PEGMAOH} = I_{PEGMA+PEGMANB} \times \frac{X_{PEGMAOH}}{X_{PEGMA+PEGMAOH}}$$

It is challenging to quantify peptide inside functional polymers using NMR because the associated peaks fall below the detection threshold of the NMR spectra. As a result, the peptide density for CHNS was estimated using trace elements microanalysis (conducted by Macquarie University Analytical and Fabrication Facility, School of Natural Sciences, NSW, Australia). In this application, bulk peptide density refers to the density of peptide in the bulk (measured in µg per mg of polymer). **Equation 1** was used to do the computation. Here, Y represents the nitrogen concentration of the polymer (measured in mg of nitrogen per kg of polymer), as determined by trace elements microanalysis. $M_{peptide}$ is the molecular weight of the peptide, $M_N$ is the molecular weight of nitrogen, n is the number of nitrogen atoms per mole of peptide (n = 12), and $10^3$ is included for unit conversion to µg/mg.

Furthermore, the number of peptides per each polymer chain (Peptides/Chain) was calculated using equation (2). In this equation, $M_{nPolymer}$ represents the polymer's number-average molecular weight as determined by GPC analysis, and $10^6$ is the unit conversion used to calculate the number of peptides per polymer chain. The results are shown in **Table 2**.

$$\text{Bulk peptide density} = \frac{Y \times M_{Peptide}}{M_N \times n \times 10^3} \qquad (1)$$

$$\text{Peptides/Chain} = \frac{Y \times M_{n\ Polymer}}{M_N \times n \times 10^6} \quad (2)$$

To create the nanocluster of multivalent ligands on the polymer surface, we followed the same approach by using a blend of non-functionalized polymer with peptide-functionalized polymer with optimized global and local densities based on our previous study [1]. We focus on RGD and AG73 peptides to provide both integrin and syndecan binding and using A2G80 which can target the dystroglycan receptor (**Figure 1 and Table 1**).

Blends of functionalized polymer (MPP, with local densities of ~4 peptides per chain) and non-functionalized polymer (MP) were prepared using appropriate blending ratios to reach global densities of 7 µg of peptide per mg of polymer mixture and dissolving the polymers with 10 wt.% in methanol:water (4:1) at 60 °C. Solution of MP and MPP alone with the same concentration and solvent mixture were also used to generate the surfaces without peptide (L0G0) and randomly distributed RGD (L1G7), respectively as control surfaces along with polymer blend with the highest global and local densities of scramble peptide (RGE/A2G80SC/AG73SC).

To generate polymer film, polymer solution was cast (50 µl.mm$^{-2}$) onto substrates (round glass coverslips, No.1, 11 mm) and then samples were slowly dried under a fume hood at room temperature for 24 h before use. For biological assays, all samples were sterilized using UV light for 2 h, 3× rinsing with sterile PBS, and then incubated with serum-free culture media at 37°C overnight before cell seeding.

#### 1.2.4. Evaluation of myoblasts' adhesion on peptide surfaces

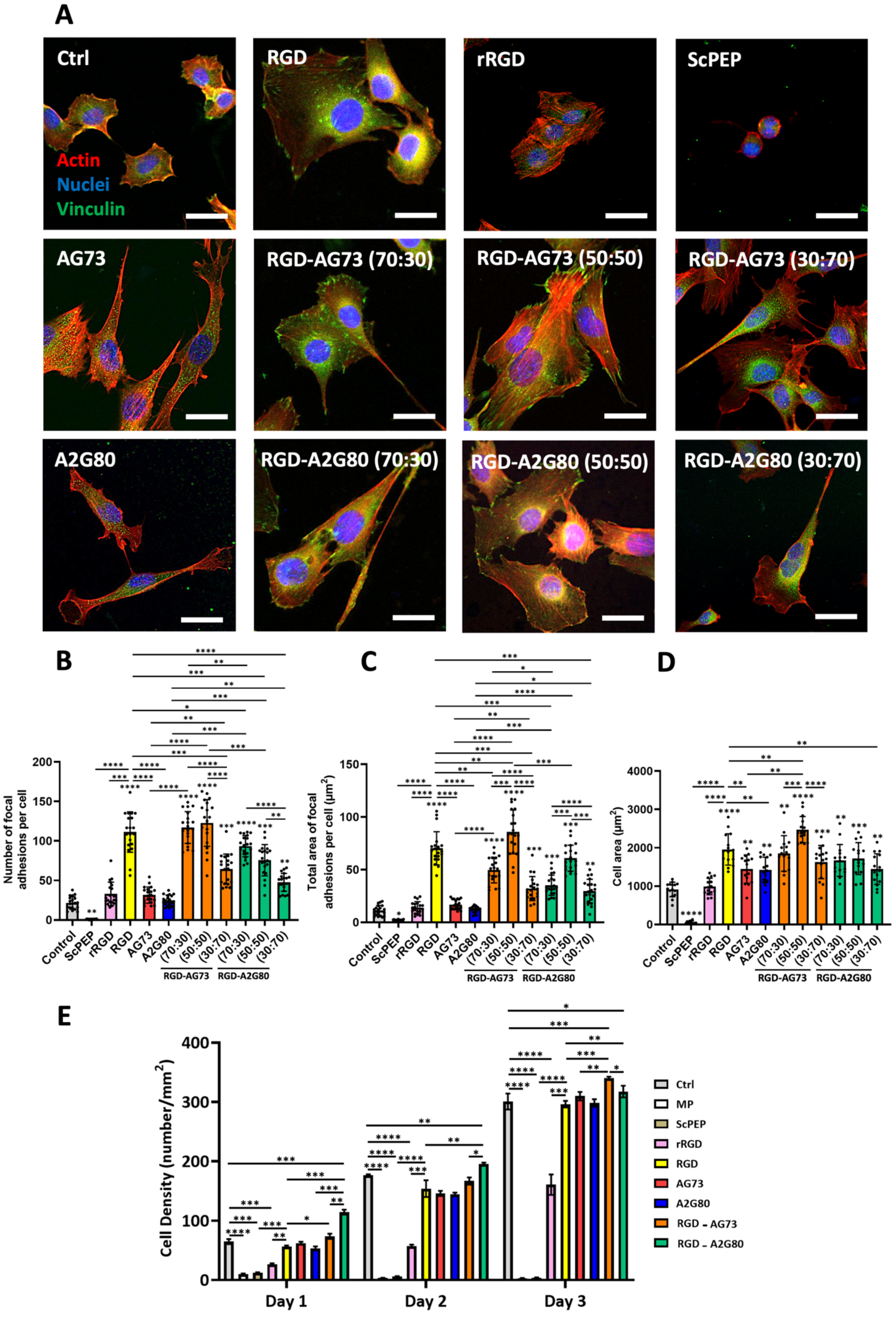

**Figure S4.** Evaluation of C2C12 myoblast proliferation with A) representative fluorescent images of cells on polymer-coated substrates with different ligand densities on day 3. Staining was done against actin filaments (Actin555, red) and cell nuclei (DAPI, blue), scale bar: 200 µm. B) Cell growth was quantified over 3 days of culture using AlamarBlue assay. ✻ Represent statistical differences between selected groups (n=9).

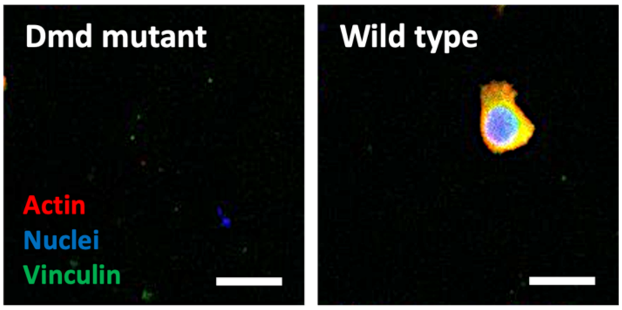


**Figure S5.** Representative confocal microscopy images of *Dmd* mutant and wild type cells cultured on the non-functionalized surface (MP) after 24 h. Staining was done against actin filaments (red), vinculin (green), and nuclei (blue). Scale bars: 30 µm.

#### 1.2.5. CRISPR-Cas9 generation of control and *Dmd* mutant C2C12 myoblast cell lines

Single guide RNAs (sgRNAs) targeting the mouse *Dmd* gene at exon 9 and 29 (sgRNA target sequence, AAGGCATAACTCTTGAATCG and CATGGATGAACTGATCAATG) and the mouse genome non-targeting Control sgRNA (AAAAAGTCCGCGATTACGTC) were purchased from Synthego (CRISPRevolution 115 sgRNA EZ Kit, Synthego, Menlo Park, CA, USA). For sgRNA/Cas9 ribonucleoprotein (RNP) formation, 0.5 µl of each *Dmd* sgRNA or 1 µl of the Control sgRNA were incubated with 4 µl of PBS (final volume of 5 µl) for 10 min at room temperature. C2C12-Cas9 myoblasts ($1.15 \times 10^5$) kindly donated by Dr. Kevin Watt (Centre for Muscle Research, Department of Anatomy and Physiology, The University of Melbourne) were washed in PBS, pelleted, resuspended in 20 µl SE Cell Line NucleofectorTM solution (#V4XC-103, Lonza, Basel, Switzerland) and mixed with the pre-prepared 5 µl sgRNA/Cas9 RNP prior to immediate electroporation using the SE Cell Line 4D-NucleofectorTM X Kit S electroporation wells (Lonza) and 4D-Nucleofector® X Unit (Lonza). 130 µl of growth media (DMEM/10% FBS/1% L-glut) was added to cells in electroporation wells, and cells were rested for 10 min at 37 ºC + 5 % $CO_2$. Cells were then transferred to one well of a 6-well plate and incubated in an additional 2 ml/well warm growth media at 37 ºC + 5 % $CO^2$, with media refreshed after 24 h. When cells reached approximately 60% confluence, they were sub-cultured in growth media and incubated at 37 ºC + 5 % $CO_2$, with media refreshed every 48 h. When cells were sufficiently confluent, they were frozen down and stored at -80°C for subsequent experiments.

#### 1.2.6. Evaluation of dystrophin expression in myotubes

To confirm knockdown of dystrophin Dp427 isoform protein expression in targeted C2C12 myotubes, control and *Dmd* mutant C2C12 myoblasts were grown to confluence over 72 hours in C2C12 GM, an induced to differentiated into myotubes via incubation in C2C12 DM for 6 days, with media refreshed every 48 h. Cells were lysed on ice in ice-cold 1× RIPA buffer (50 mM Tris-HCl (pH 7.4), 150 mM NaCl, 0.25% deoxycholic acid, 1% NP-40, 1 mM EDTA) supplemented with protease inhibitor cocktail (1:1000) and 1 mM PMSF. Lysates were prepared by scraping with a pipette tip, sonicated (15 s, Microson XL-2000), and centrifuged (10,000 rpm, 4 °C, 10 min) as described previously (REF YOUR PREVIOUS PAPER HERE AS WELL AS THAKUR SS BIOLOGY OPEN 2020). Total protein concentration was assessed with a DC protein assay (Bio-Rad Laboratories), and samples were mixed with Laemmli buffer [4×; 0.25 M TrisHCl (pH 6.8), 6% SDS, 40% glycerol, 0.04% bromophenol blue, 16% Dithiothreitol (DTT)] at a concentration of 2 mg/ml and heated to 95 °C for 3 mins.

Samples were run on 4–20% Criterion TGX Stain-Free gels (Bio-Rad Laboratories) at 100–160 V for 60 min at room temperature. Proteins were transferred to PVDF membranes (1 h at 300 mA), visualized for total protein using a Chemidoc MP system, blocked with 5% BSA in TBST, and incubated overnight at 4 °C with primary antibodies (mouse-α-dystrophin MANEX1011B(1C7) (1:1000, AB-1157876) and mouse-$\alpha$-MF20 (1:1000), Developmental Studies Hybridoma Bank, The University of Iowa; USA) and mouse-β-dystroglycan (1:100; B-DG-CE; Leica Biosystems)). After washing with Tris-buffered saline-Tween 20 (TBST), membranes were incubated for 1 h in HRP-conjugated sheep-α-mouse IgG in 5% BSA/TBST (1:5000, GE Life Science), washed with TBST, and reacted with either Immobilon Forte (for MF20 and β-dystroglycan detection; Merck Millipore) or SuperSignal West Femto (for dystrophin detection; Thermo Fisher Scientific_ enhanced chemiluminescence reagent).

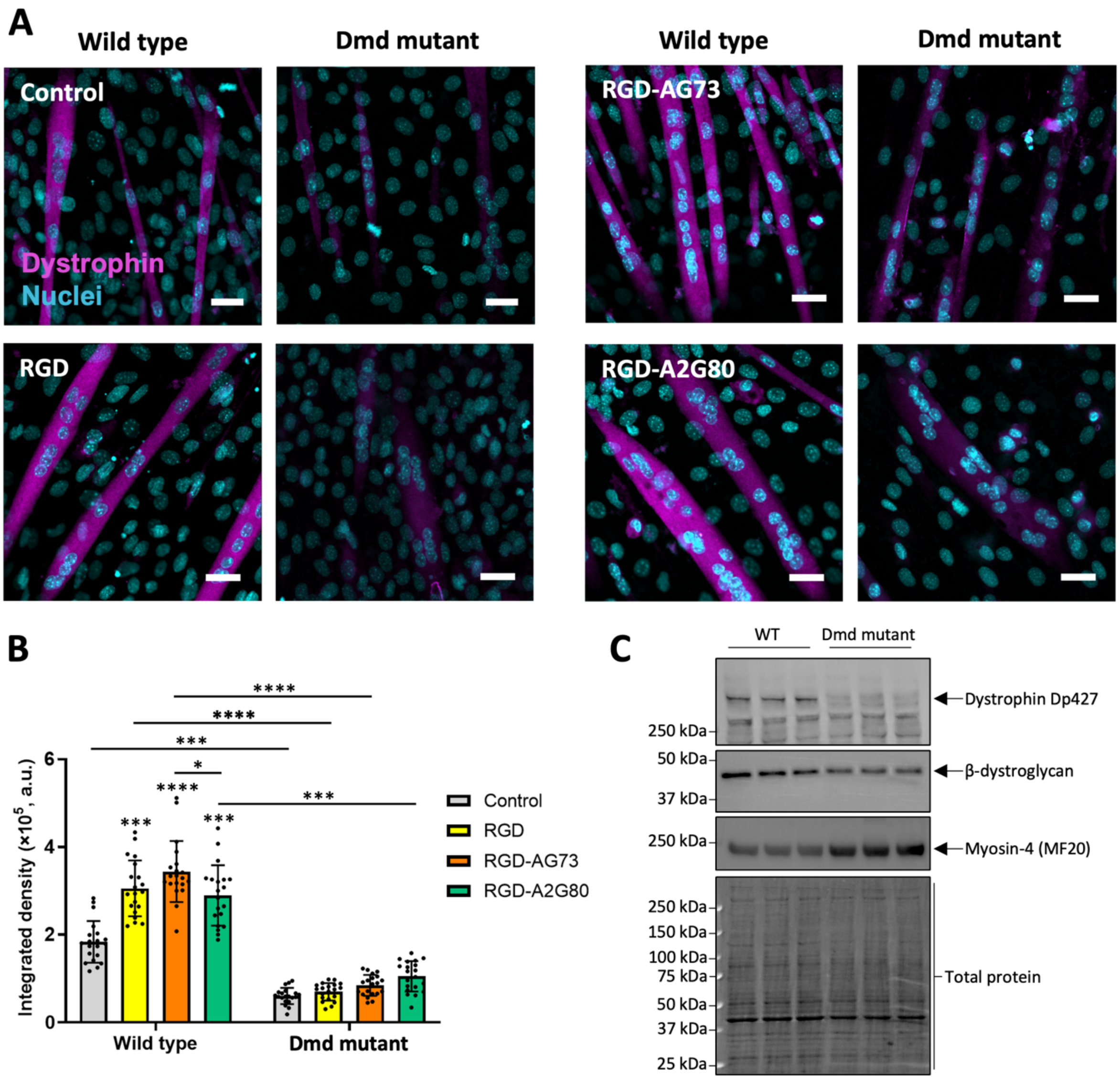


**Figure S6.** A) Confocal imaging reveals the dystrophin deficiency in *Dmd* mutant cells on all surfaces after 7 days. Staining against dystrophin (DAG1, magenta) and nuclei (cyan), scale bar: 30 µm with B) calculated fluorescent signal density. (C) Western immunoblotting confirmed loss of Dp427 protein expression and lower β-dystroglycan protein expression in *Dmd* mutant C2C12 myotubes relative to wild type control myotubes. ✻ Represent statistical significance. Asterisks directly above data points indicate statistical differences with control and asterisks above the bars indicate statistical differences between treatment groups.

#### 1.2.7. Evaluation of motor neurons' adhesion on peptide surfaces

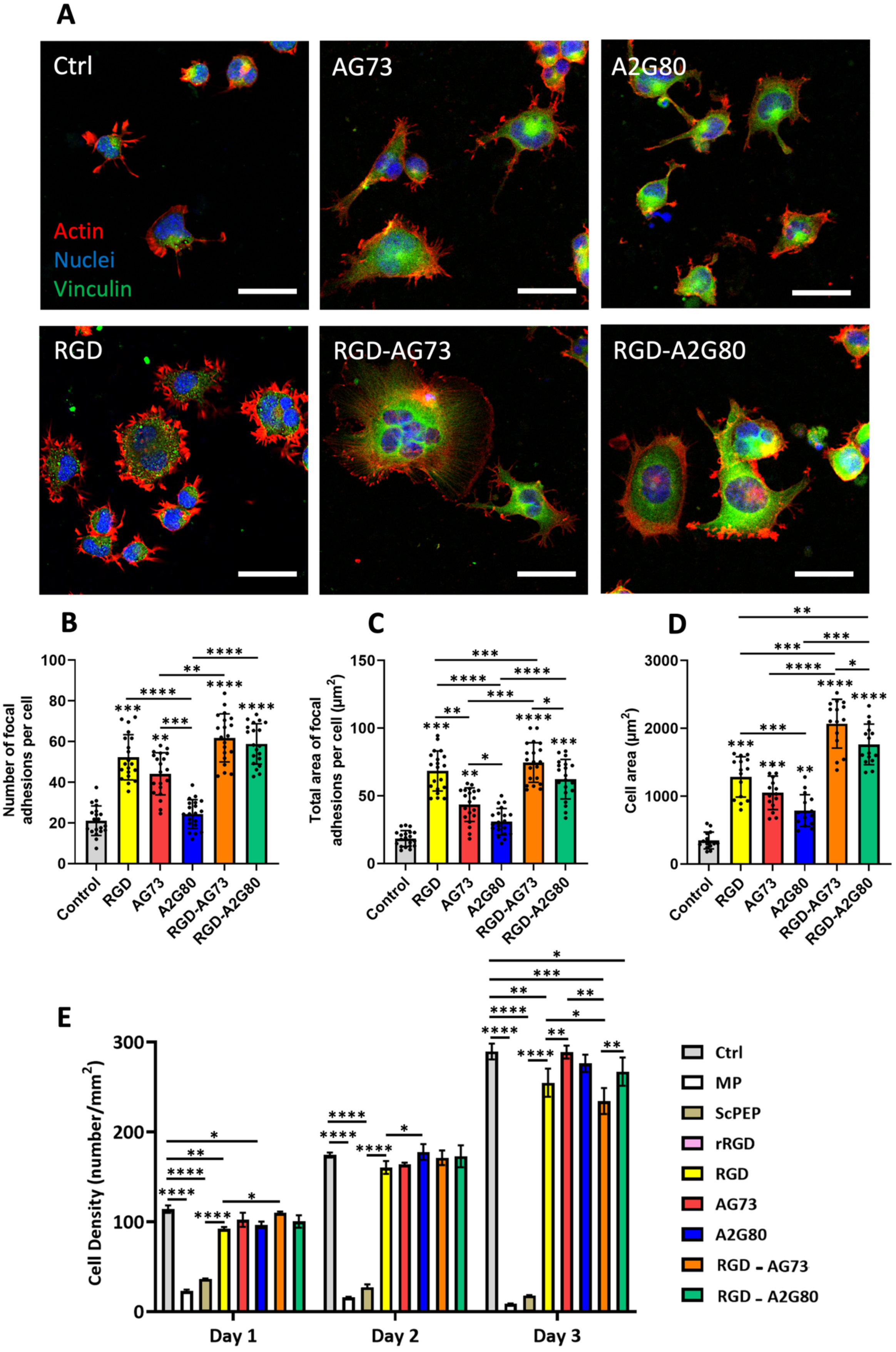

**Figure S7.** Evaluation of motor neuron proliferation with A) representative fluorescent images of cells on polymer-coated substrates with different ligand densities on day 3. Staining was done against actin filaments (Actin555, red) and cell nuclei (DAPI, blue), scale bar: 200 µm. B) Cell growth was quantified over 3 days of culture using AlamarBlue assay. ✻ Represent statistical differences between selected groups (n=9).

#### 1.2.8. Evaluation of actin cytoskeleton and receptor localization in myotubes

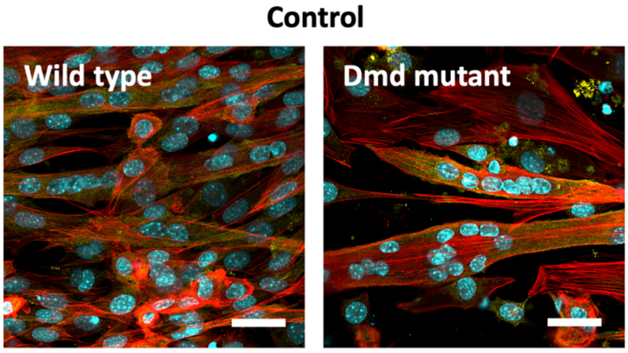


**Figure S8.** Representative confocal microscopy images of cells cultured on control surface after 7 days. Staining was done against actin filaments (red), dystroglycan (yellow), and nuclei (cyan), Scale bars: 30 µm.

#### 1.2.9. Evaluation of neuromuscular junction formation

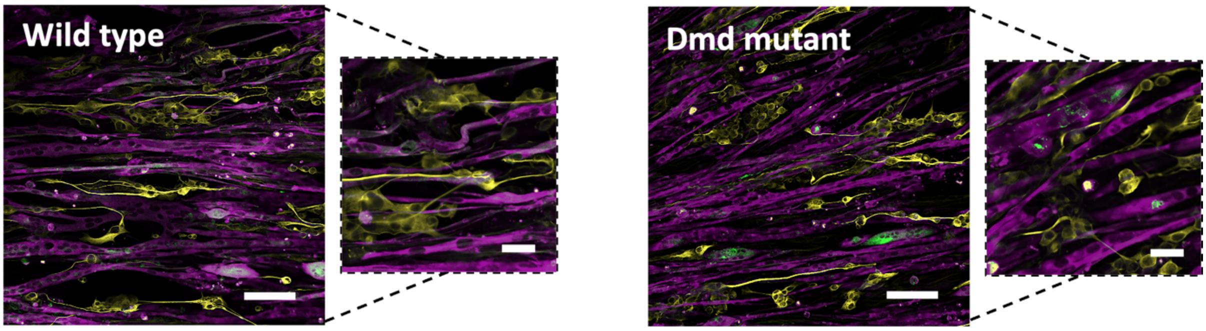


**Figure S9.** Representative confocal images of neuromuscular connectivity after 14 days of culture on RGD homocluster surface in *Dmd* mutant and wild type cells. Staining was performed using MyHC (MF20, magenta), AChR (α-BTX, green), and β-tubulin III (yellow). Scale bars: 100 µm and 30 µm for the zoom-in panel.
